# Three million images and morphological profiles of cells treated with matched chemical and genetic perturbations

**DOI:** 10.1101/2022.01.05.475090

**Authors:** Srinivas Niranj Chandrasekaran, Beth A. Cimini, Amy Goodale, Lisa Miller, Maria Kost-Alimova, Nasim Jamali, John G. Doench, Briana Fritchman, Adam Skepner, Michelle Melanson, Alexandr A. Kalinin, John Arevalo, Marzieh Haghighi, Juan Caicedo, Daniel Kuhn, Desiree Hernandez, Jim Berstler, Hamdah Shafqat-Abbasi, David Root, Susanne E. Swalley, Sakshi Garg, Shantanu Singh, Anne E. Carpenter

## Abstract

Identifying genetic and chemical perturbations with similar impacts on cell morphology can reveal compounds’ mechanisms of action or novel regulators of genetic pathways. Research on methods for identifying such similarities has lagged due to a lack of carefully designed and well-annotated image sets of cells treated with chemical and genetic perturbations. Here, we create such a Resource dataset, CPJUMP1, where each perturbed gene is a known target of at least two chemical compounds in the dataset. We systematically explore the directionality of correlations among perturbations that target the same gene, and we find that identifying matches between chemical perturbations and genetic perturbations is a challenging task. Our dataset and baseline analyses provide a benchmark for evaluating methods that measure perturbation similarities and impact, and more generally, learn effective representations of cellular state from microscopy images. Such advancements would accelerate the applications of image-based profiling, such as functional genomics and drug discovery.

## Introduction

Image-based profiling of cell samples is proving increasingly useful for biological discovery ^1^. In image-based profiling, cells are treated with perturbations of interest and the resulting morphology is captured by microscopy. Cell morphology is quantitatively compared to identify meaningful similarities and differences among the perturbations, in the same way that transcriptional profiles are used to compare samples. More than a dozen applications have been demonstrated ^1^, including: (a) identifying the mechanisms of a disease by comparing cells from patients with a disease to healthy patients, (b) identifying the impact of a chemical compound by comparing cells treated with it to untreated cells, and (c) identifying gene functions by clustering large sets of genetically perturbed samples to determine relationships among the genes.

Typically, morphological features are extracted from each cell using classical image processing software. These so-called “hand-engineered” features have been carefully developed and optimized to capture cellular morphology variations, including size, shape, intensity, and texture of the various stains in the image. These features are the current standard in the field and require post-processing steps including normalization, feature selection and dimensionality reduction ^2^. With advances in representation learning during the last decade, it is natural to ask what set of features could be automatically identified by machine learning algorithms, directly from pixels.

However, image-based profiling has yet to fully benefit from the latest machine learning research. The vast majority of studies use classical segmentation and feature extraction; deep learning methods are beginning to be explored and there is much room for advancement ^3,4^. Historically, the lack of ground truth has been a major limiting factor in the field, that is, the “correct” relationships among perturbations (e.g., genes and compounds) are unknown and require significant effort to ascertain ^5^. While this is exciting because the potential for biological discovery is high, the lack of ground truth presents a challenge for optimizing deep learning pipelines.

Here, we describe our design and creation of a benchmark dataset via a single large experiment, CPJUMP1. The dataset comprises roughly three million images of cells, image-based profiles of seventy five million single cells, and well-level aggregated profiles. A sample five channel image is shown in Figure 1. This dataset contains chemical and genetic perturbation pairs that target the same genes and are tested in separate wells to see whether they produce similar (or opposite) phenotypes. Although these pairings are not absolute “truth” for a number of reasons discussed later, they are nevertheless more likely than random pairs to match (i.e., induce similar/opposite morphology changes). This Resource is unique because there are no other image-based datasets of the Cell Painting assay (described later) that include pairs of annotated genetic and chemical perturbations performed side by side, under different experimental conditions such as different cell types, time points, and imaging conditions. These were also executed in parallel to minimize technical variations that may confound the signal.

**Figure 1:**
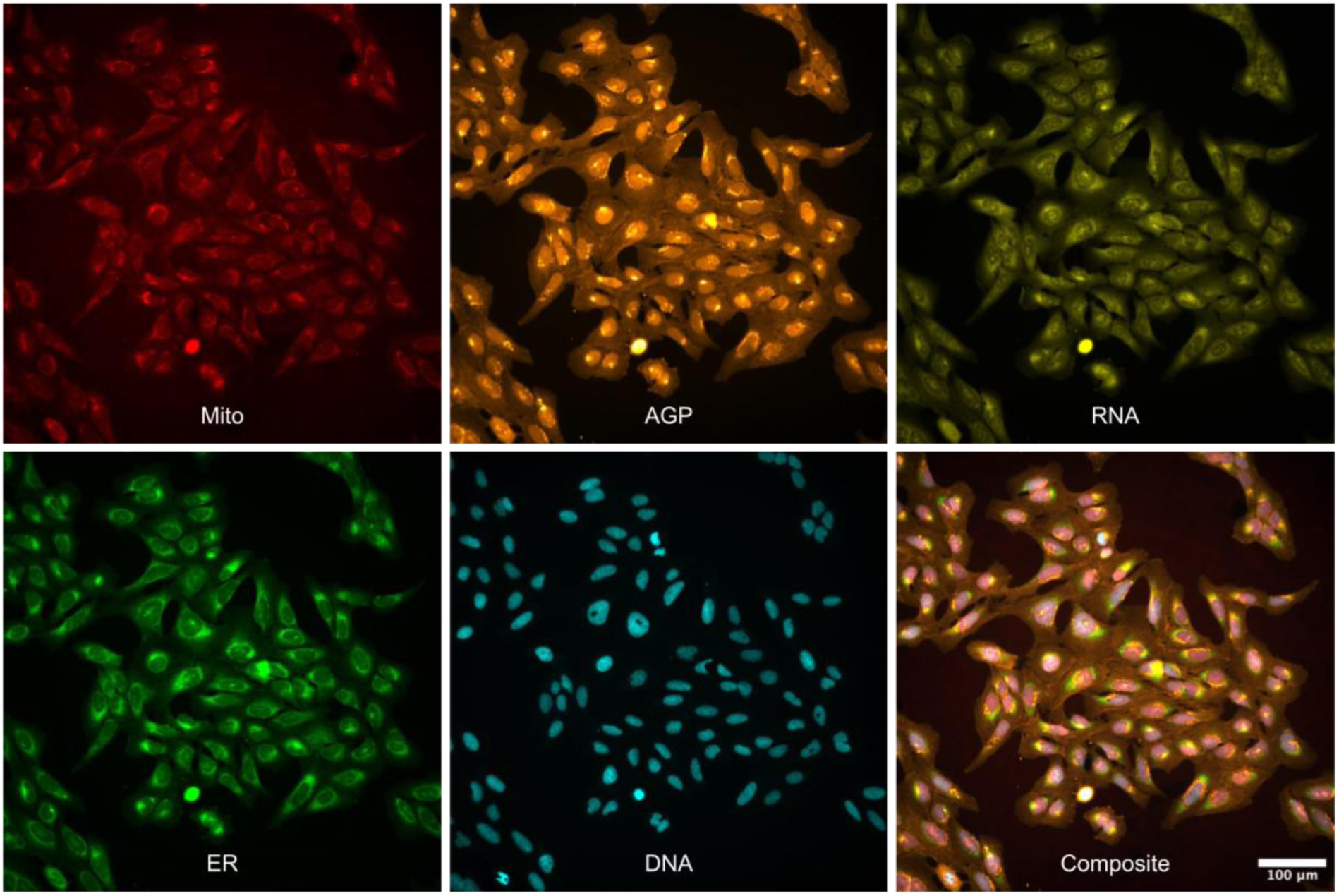
Sample images from the dataset. A 5-channel image of human U2OS cells treated with the compound PFI-1. The channel names indicate the cellular structures identified in each image (see online methods for details; AGP = Actin, Golgi, Plasma membrane). Other example images (including brightfield channels not shown here) are available at https://github.com/jump-cellpainting/2023_Chandrasekaran_submitted/tree/5c6fcf9dc70a85176f5afc5263acbc230d90ca40/example_images. Scale bar = 100 µm.

There are many public Cell Painting datasets (for example, https://github.com/broadinstitute/cellpainting-gallery) but we are only aware of one with genetic and chemical perturbation types run in parallel (RxRx3, https://www.rxrx.ai/rxrx3), it has not been provided with gene-compound relationship annotations; any that exist would be scattered across many plates and batches. As well, it includes only a single cell type, time point, and imaging condition. Thus, the nature of this dataset allows testing computational strategies to optimally represent the samples so that they can be compared and thus uncover valuable biological relationships. It also allows comparison of CRISPR-Cas9 knockout and ORF (Open Reading Frame) overexpression as mechanisms to perturb genetic pathways and to identify compounds’ mechanisms of action.

## Results

To push forward advancements in this field, we assembled a consortium of ten pharmaceutical companies, two non-profit institutions, and several supporting companies, known as the JUMP-Cell Painting Consortium (Joint Undertaking in Morphological Profiling). Members of this Consortium created the ground truth dataset we present here, for optimizing the main assay used in image-based profiling, called the Cell Painting assay, and to move methods in the field forward ^6,7^. We selected and curated a set of 160 genes and 303 compounds with (relatively) known relationships among each other, and designed an experimental layout to enable testing and comparing methods to quantify their similarities (online methods), all with a strong emphasis on making this dataset useful for developing computational methods for the field.

There are two groups of experimental conditions in this dataset, the primary group and the secondary group. In the primary group, we separately captured chemical and genetic perturbation (CRISPR knockout and ORF overexpression) profiles in two cell types (U2OS and A549) at two time points (Supplementary Table 1; a representation of profiles from a subset of this experiment is shown in Figure 2). There are 40 384-well plates in the primary group of experimental conditions (Figure 3). The secondary group consists of additional plates of experimental conditions as well as plates from the primary group subjected to additional imaging conditions (Figure 3), which are described in online methods. In addition to being used for optimizing the assay conditions ^7^, the primary and secondary groups offer multiple “views” of cells treated with each chemical or genetic perturbation, and therefore can be used for many interesting machine learning explorations, such as style transfer e.g., to attempt prediction of one experimental condition from another, information retrieval and multi-view learning, and benchmarking representation learning methods. There are 11 additional plates in the secondary group, but there are 67 plates of images because several plates were imaged multiple times (Figure 3; online methods).

**Figure 2:**
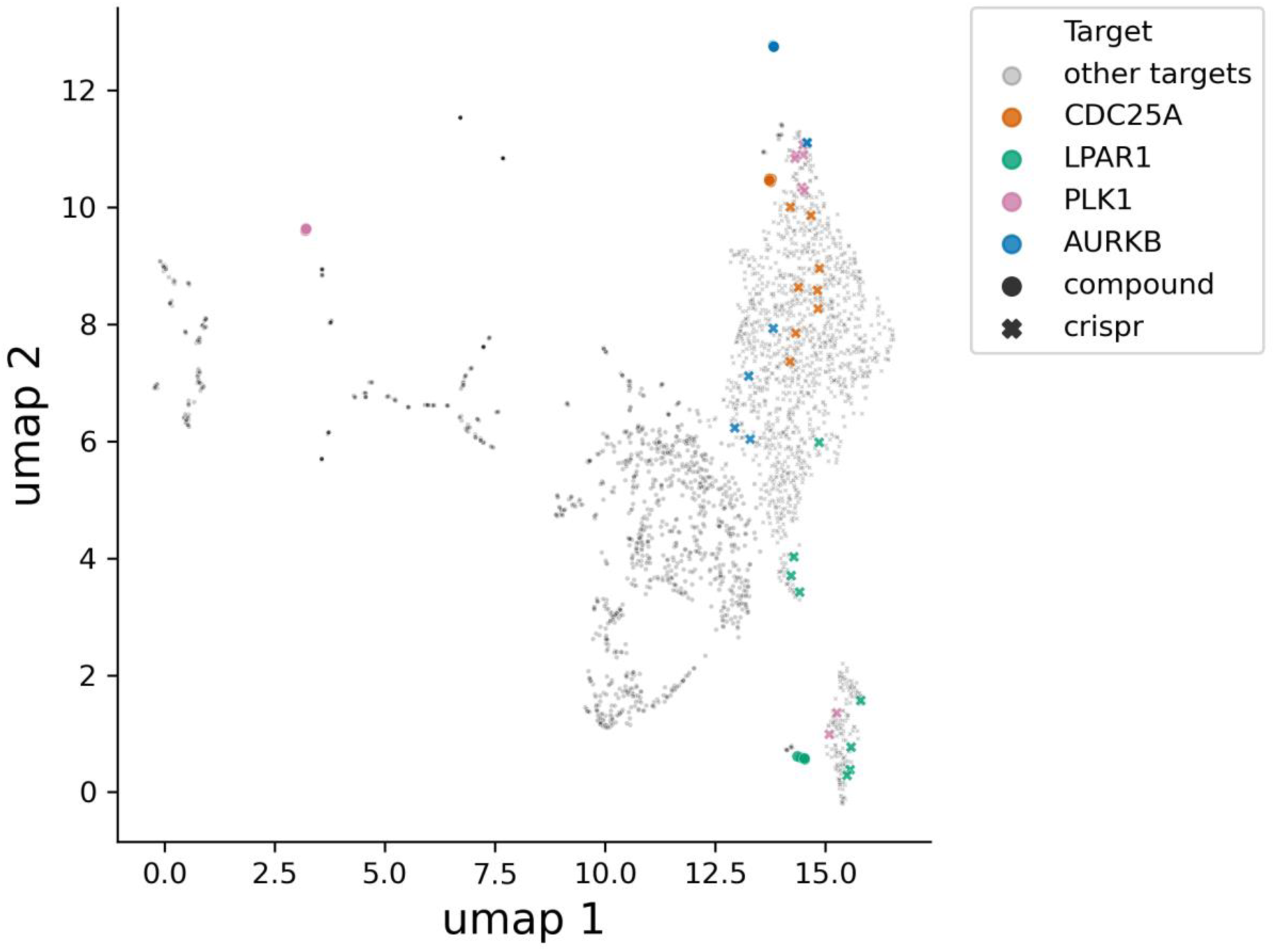
UMAP representation of image-based profiles from a subset of the primary group of tested samples. Profiles from human A549 cells perturbed by all compounds and CRISPR guides at the long time points (Supplementary Table 1) are shown here. The top four most-similar compound-genetic perturbation pairs, that target the same gene, are highlighted.

**Figure 3:**
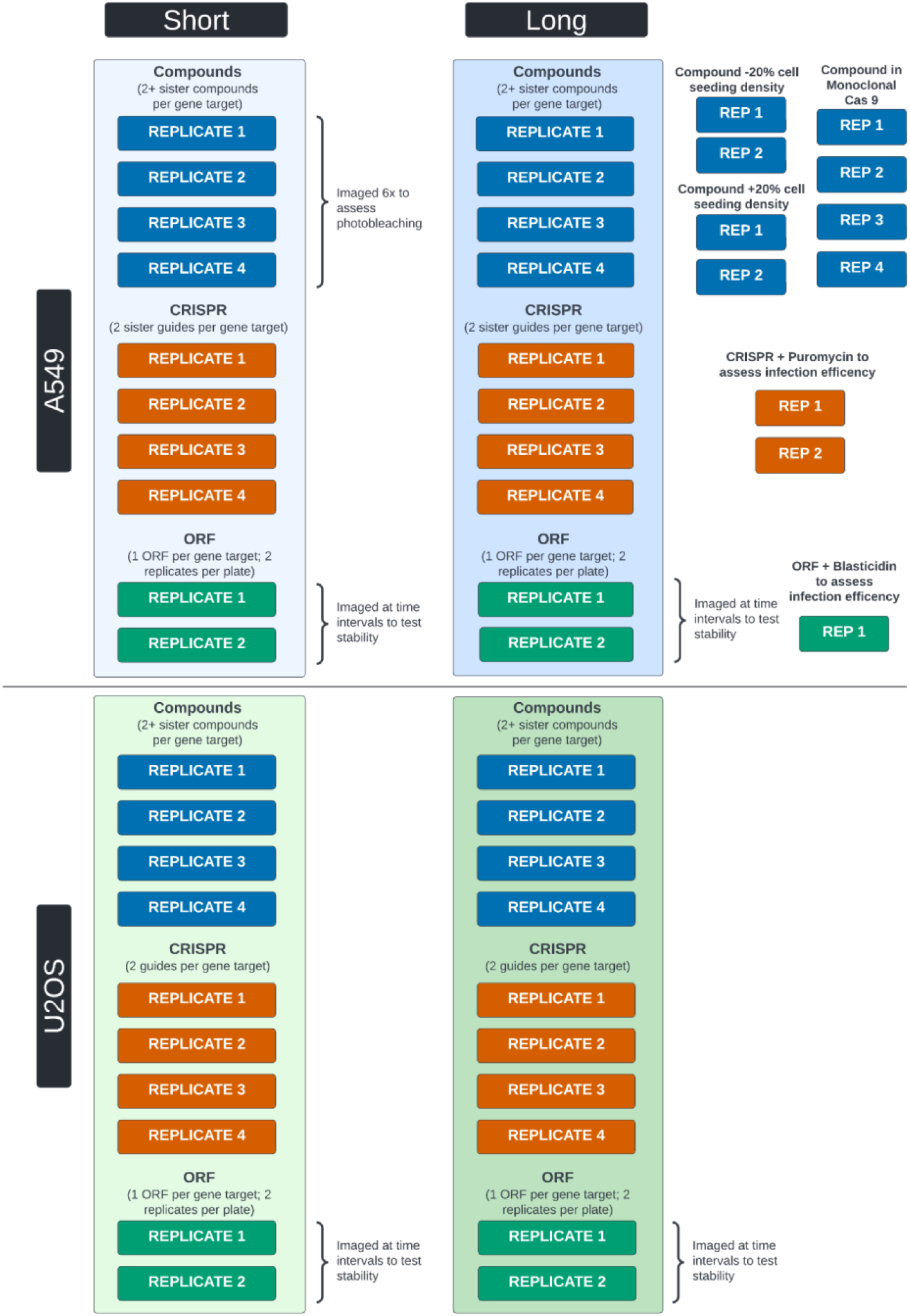
Schema of all the data generated in CPJUMP1 experiment. Each rectangular block of replicates (REPLICATE X or REP X) is a 384-well plate of cells perturbed by compounds, CRISPR guides or ORFs and subjected to different experimental conditions. Short and long time points are described in Supplementary Table 1. Plates in the vertical blue and green boxes comprise the primary CPJUMP1 experiment. The other experimental conditions are further described in the online methods.

A major goal of image-based profiling is to derive a representation from the quantitative images of cell samples, such that samples in biologically similar states have similar representations. Given such a representation, solutions for many of the applications discussed become immediately feasible. Here, we benchmark representation learning as a foundation for methods development in the future.

As a way to compare different representation methods, we created benchmarks based on two tasks: (1) detecting differences between perturbations and negative controls to identify “active” perturbations, and (2) grouping gene-compound pairs where the gene is a target of the compound (as well as grouping two CRISPR guides targeting the same gene, or two compounds annotated as targeting the same gene). For both tasks, we use cosine similarity (or its absolute value) – a simple but widely used correlation-like metric – to measure similarities between pairs of well-level aggregated profiles. In some cases, the expected directionality of correlation is positive while in other tasks, correlations may be strongly positive or negative; we adjust statistical tests for each task accordingly.

### Benchmarking representation learning methods: perturbation detection

We chose perturbation detection as one of the tasks to evaluate representations because it often precedes other useful applications (by removing samples that have no/little true signal), and is equivalent to measuring statistical significance of the perturbation’s signal. For example, a set of chemical or genetic perturbations might be filtered by this criterion before embarking on subsequent laboratory experiments, or prior to training a model, or other analysis that could be confounded by noisy signals. It can also be useful for determining what experimental protocol or computational analysis pipeline is most sensitive among several alternatives. It should be noted that even given perfect computational methods for feature extraction, batch correction, and profile comparison, many samples will be detectably different from negative controls for several biological reasons. For example, a chemical or genetic perturbation may only impact cell morphology in a particular cell type, under particular environmental conditions, at a particular time, or if particular stains were used, conditions which may not have been met in the experiment. Conversely, a perturbation’s impact may be amplified by the plate layout, as even unrelated perturbations in the same well position might look similar. This concern is overcome by matching treatments in different well positions, where such data is available (see Benchmarking perturbation matching methods).

We used *Average Precision (AP)* to measure each primary group sample’s ability to retrieve its replicates against the background of negative control samples, using cosine similarity as the similarity metric. The significance of the AP value is assessed using permutation testing to obtain a p-value, which is then FDR adjusted to yield a q value. We average AP for each perturbation to obtain mAP and then call the fraction of perturbations with a q value below the significance threshold (0.05) *Fraction Retrieved (FR)*. Details about the computation of AP and FR are provided in the online methods.

In general, we find that FR for compounds is higher than that of genetic perturbations, across all conditions (Figure 4a). This indicates that chemical compounds produce phenotypes that are more distinguishable from negative controls, as compared to phenotypes produced by CRISPR knockout and ORF overexpression. We also find that FR is higher for CRISPR knockout than for ORF overexpression (Figure 4a). In summary, compounds, CRISPRs, and ORFs all yield signals in the assay, with the compounds being the strongest and ORFs the weakest. However, we emphasize a strong technical variable that precludes a strong conclusion here: the reduced FR values for ORF may be attributed to plate layout effects, where identical treatments in different rows or columns have dissimilar profiles. This factor amplifies the systematic technical noise in the compound and CRISPR plates due to their particular layout, while it adversely impacts ORFs (online methods). Retrieving the same position replicates for ORF does (likely artificially) increase FR (Supplementary Figure 1). Plate layout effects may be partly mitigated by mean centering every feature at each well position, though we have not applied the correction to this dataset. Furthermore, though there is a presence of signal for all three perturbation modalities, we note that these were not random sets of genes and compounds; instead, the compounds were chosen from the Drug Repurposing set ^8^ and the genes as being targeted by those compounds. This likely enriches for reagents that yield phenotypes versus random genes and compounds.

**Figure 4:**
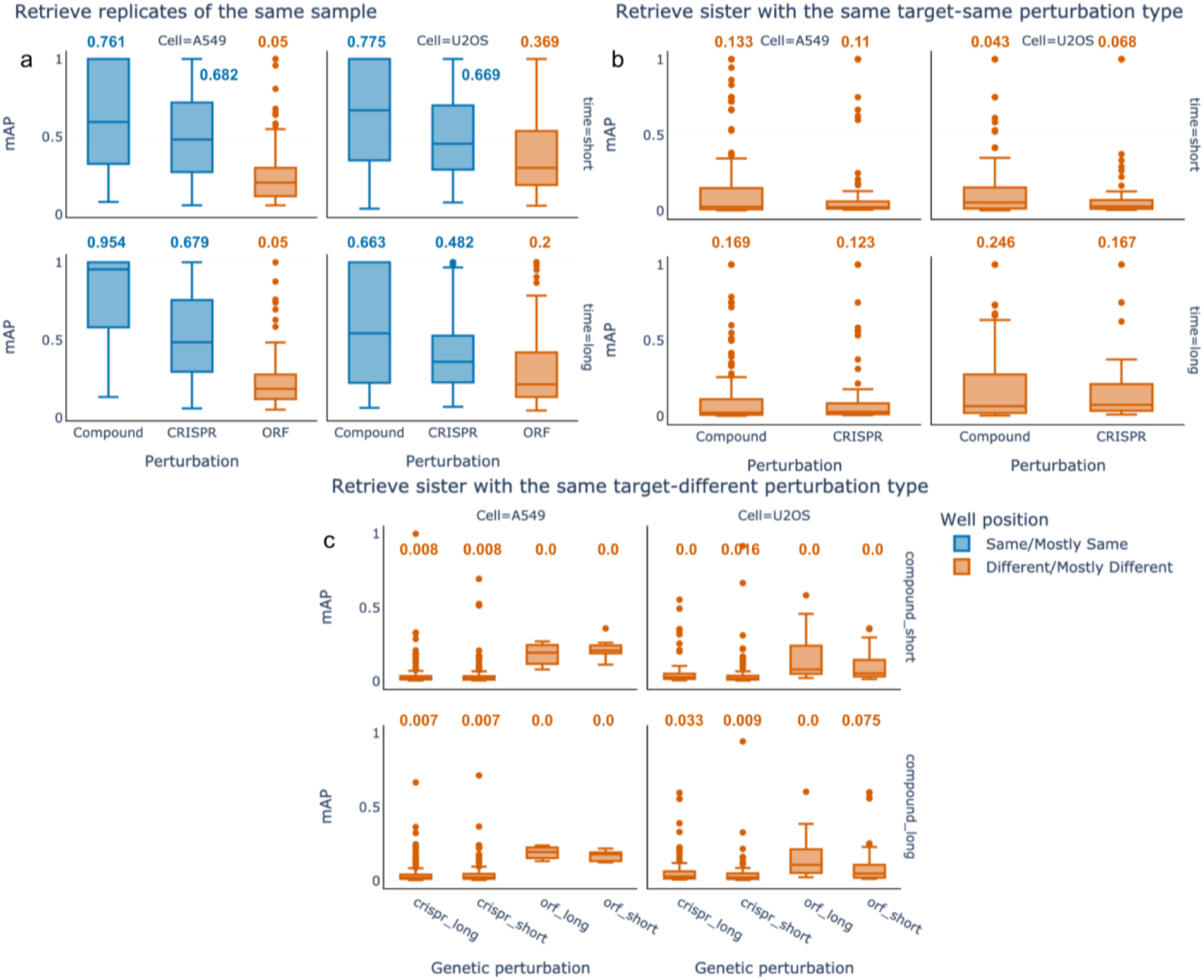
Benchmark results for progressively more difficult retrieval tasks. a) Perturbation detection: mAP for perturbation detection is shown across experimental conditions – cell type (columns) and time points (rows; short and long time points are defined in Supplementary Table 1). The numerical values shown above each box plot are the fraction of perturbations that can be successfully retrieved (FR) values for each retrieval task. Box plot boundaries are 75th (Q3) and 25th (Q1) percentiles, with whiskers at +/- 1.5 times the interquartile range (Q3-Q1) or the highest or lowest data point. The colors of the bars denote whether the query perturbation and retrieved perturbation are in mostly the same well position (blue) or mostly different well positions (red); the latter is a more challenging task due to technical well-position artifacts. b) Perturbation matching, within a perturbation type: the plot shows mAP for sister perturbation retrieval (that is, pairs of compounds or pairs of CRISPR guides annotated with the same gene target). ORFs are not shown because there is only a single ORF reagent per gene target. Absolute cosine similarity is used for calculating mAP values for compounds because pairs of compounds annotated to target the same gene can be positively or negatively correlated. c) Perturbation matching, across perturbation types: the plot shows mAP values for retrieving compound-gene pairs. Absolute cosine similarity is used for calculating mAP values for both compound-CRISPR and compound-ORF matching. The number of data points in each box plot is available in Supplementary Table 2

### Benchmarking representation learning methods: perturbation matching

We next established a benchmark for researchers to develop and test strategies for a real-world retrieval task, where we search for genes or compounds that have a similar impact on cell morphologies as the query gene or compound. Improved methods would allow for improved discovery of compounds’ mechanisms of action based on a compound query ^9^ and virtual screening for useful compounds based on a gene query ^10^. This dataset presents a unique opportunity to match profiles of perturbations across modalities (chemical versus genetic) because genes in this dataset are targeted by two types of genetic perturbations (ORF and CRISPR-Cas9 knockout) and by at least two compounds. Similarly, because there are both a pair of CRISPR guides and a pair of compounds targeting each gene, this dataset can be used for matching profiles within a perturbation modality (there is only one ORF reagent per gene, so a similar analysis is not possible for overexpression). It also offers an opportunity to study the directionality of profile matching; for example, whether CRISPR knockouts and ORF overexpressions consistently yield anti-correlated profiles.

After filtering out perturbations that were indistinguishable from negative controls (q value>0.05), we then evaluated AP to identify “true” connections (perturbation pairs that target the same gene) that are distinct from “false” connections (pairs not known to target the same gene).

We first tested the ability to retrieve true connections *within* the same perturbation modality – that is, “sister” compounds that are annotated as targeting the same gene should match each other, and “sister” CRISPR guides that target the same gene should match each other. Because compounds can enhance or inhibit the function of a gene (or have other impacts), those with the same gene annotation might be positively correlated or negatively correlated; for compounds, we therefore used the absolute value of cosine similarities while calculating AP. By contrast, for CRISPR guides, those targeting the same gene are expected to be positively correlated, so we used the actual values of cosine similarities, as in the perturbation detection task.

With baseline methods, this task is (not surprisingly) much more challenging than retrieving replicates of the same sample (compare Figure 4a and 4b). Of all compounds that yield a signal, only about 5-25% of them correctly match their sister compounds targeting the same gene. Likewise, and more surprisingly given their expected accurate annotation and specificity, only 7-17% of the CRISPR reagents correctly match to their sister guides targeting the same gene. We cannot distinguish the many factors that likely make this a challenging task, including non-optimal ground truth annotations for compounds, off target effects (for compounds and CRISPR guides), differing levels of knockdown for CRISPR guides, lack of information content in the assay, polypharmacology where each compound impacts multiple targets (see Discussion) and/or non-optimal methods for matching samples. Though the FR values are similar for compounds and CRISPR guides, more compounds are distinguishable from negative control compared to CRISPR guides (Figure 4a, Supplementary Table 2). Thus, surprisingly, retrieving sister compounds is more successful than retrieving sister CRISPR guides (Figure 4b), perhaps because compounds tend to induce stronger phenotypes (Figure 4a).

Next, we assessed cross-modality matching; that is, the ability to retrieve correct gene-compound pairs. Retrieving compound-gene pairs is more difficult than perturbation detection and sister perturbation retrieval (which itself reached only ∼25% in the best scenario), but is extraordinarily useful for identifying novel chemical regulators of genes and identifying the mechanism of query compounds. Even a low success rate, therefore, can accelerate drug discovery. Given the potential for compound-gene matches to be positive or negatively correlated (detailed in the next paragraph), we used the absolute cosine similarities for both compound-ORF and compound-CRISPR retrieval.

Gene-compound retrieval is only slightly better than expected by chance (Figure 4c), and, as expected, less effective than compound-compound and CRISPR-CRISPR matching (Figure 4b). This might reflect that gene-compound annotations are less reliable than compound-compound annotations (and certainly CRISPR guide annotations, where the target gene is designed a priori to be accurate and specific), and/or that better methods are needed to align data across modalities. Comparing the two genetic perturbation modalities, we find that mAP values for retrieving compound-CRISPR pairs were better than that of compound-ORF pairs across various cell types and timepoints, except at one time point where ORFs match better to compounds than CRISPR reagents do (Figure 4c).

Although the performance for compound-genetic perturbation retrieval is low compared to the other retrieval tasks discussed above, it should be strongly noted that significant time and resources are otherwise required to identify the target of a compound, and similarly to identify compounds that target a particular gene. Therefore, even the baseline’s relatively low matching rates might accelerate drug development by yielding a list of possibilities for biologists to test directly in subsequent experiments, or to be combined with orthogonal lists of candidate target genes for a compound, to improve accuracy. Improving image representations and therefore the accuracy of predicted matches by a few percentage points could therefore have a major impact on the discovery of compounds that impact the function of genes of interest, and the identification of the mechanism of action of compounds of interest.

Finally, we examined the directionality of gene-compound matching, that is, whether a compound targeting a given gene has a correlating or anti-correlating profile with CRISPR (which reduces the amount of gene product) and ORF (which increases the amount of gene product), respectively. Most compounds are annotated as inhibiting the function of their target gene’s product, so one might expect image-based profiles from cells treated with CRISPR guides to generally positively correlate to (mimic) the corresponding compound’s profile, whereas ORF profiles might generally be expected to anti-correlate (oppose) the corresponding small molecule’s profile because overexpression often increases a gene’s function. By the same rationale, ORFs and CRISPR guides targeting the same gene might be expected to yield opposite (anti-correlated) effects on the cells’ profiles. However, we strongly note that there will be numerous exceptions given the non-linear behavior of many biological systems and any number of distinct mechanisms by which these general principles may not hold, which we have previously detailed ^10^. For example, many compounds do not inhibit their target gene’s function but instead activate it or induce some new function, and many overexpressed genes may have no impact at all, or even have a dominant negative or feedback loop/compensatory impact on the gene’s function. Furthermore, the choice of cell type, time point or readout for capturing the similarity may not be optimal. In fact, the exceptions may be more common than the common-sense rules in this case. One aim of generating this dataset was to quantify how often the expected relationships and directionalities occur, to provide concrete evidence in this so-far theoretical debate with only anecdotal evidence available.

We began by testing the basic hypothesis that CRISPR and ORF reagents targeting the same gene should yield negatively correlated (opposite) profiles to each other. Surprisingly, we found that the CRISPR and ORF profiles are slightly positively correlated with each other, in both cell types (Figure 5a, Supplementary Table 3). We then compared the cosine similarities between compound-CRISPR pairs and compound-ORF pairs that target the same gene. In both U2OS and A549 cells, we found that both compound-CRISPR and compound-ORF pairs were more positively correlated than negatively correlated (Figure 5b, Supplementary Table 4).

**Figure 5:**
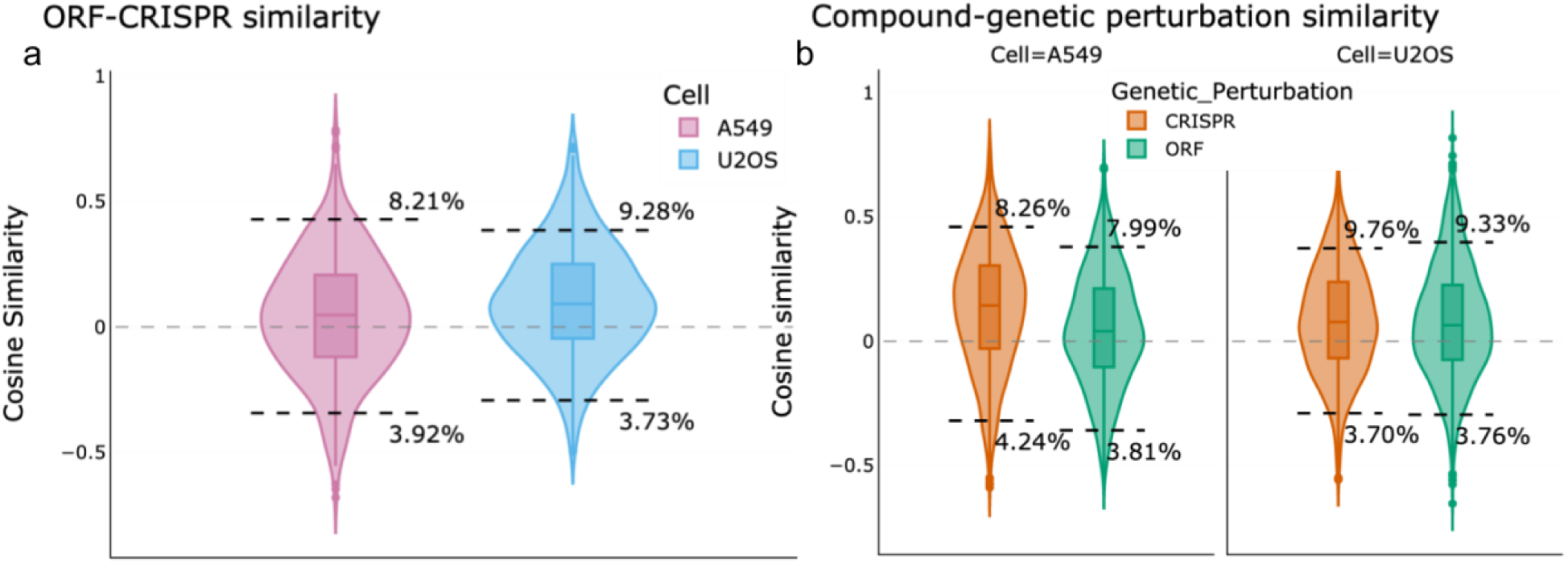
Directionality of matching cross-modality. a) Cosine similarity between ORF and CRISPR reagents targeting the same gene are shown for the two cell types, A549 (pink) and U2OS (blue). The 5th and 95th percentile thresholds of their respective nulls (ORFs and CRISPR reagents targeting different genes), along with the percentage of true pairs beyond the thresholds, are shown. We performed Fisher’s exact test to determine whether the true pairs beyond the threshold are more likely to be positively correlated or negatively correlated. We find them to be significantly more positively correlated (p values are available in Supplementary Table) 3). b) Cosine similarity between compound and the two genetic perturbation modalities, CRISPR (orange) and ORF (green) targeting the same genes. All analyses here were also statistically significantly more positively correlated; p values are available in Supplementary Table 4. Cosine similarity of zero is shown as a gray dashed line in both subplots. There are 3728 data points in each violin plot in a) and 1864 data points in each violin plot in b).

Looking at the strongest-matching positive and negative gene-compound pairings (Supplementary Tables 5 and 6), we found many pairings with explainable directionality (e.g., CRISPR knockout of a gene matches a compound annotated as that gene’s inhibitor). For example, the top positively correlated gene-compound match in U2OS cells is the PLK1 inhibitor compound BI-2536 matched with CRISPR against PLK1, and the next two matches are annotated as Aurora kinase inhibitors that match CRISPR against AURKB (Supplementary Table 5). Similarly, the five strongest matches in A549 cells (Supplementary Table 6) are all CRISPR reagents positively correlating with compounds annotated as targeting the correct gene. Some overexpressions also match the expected directionality, such as the second strongest negatively correlated match in U2OS cells, which is the compound GSK2110183, an AKT inhibitor negatively correlated to overexpression of AKT1. Still, there were many pairings with unexpected directionality, possibly due to ORFs exhibiting dominant-negative behavior, as in the seventh positively correlated gene-compound match in U2OS cells, the compound pentostatin, which is annotated as an adenosine deaminase inhibitor; it matches with overexpression of ADA (adenosine deaminase). Some CRISPR reagents produce surprising directionality, as in the case of the compound TG-003 in U2OS cells, annotated as a CLK inhibitor but negatively correlated to CRISPR knockout of CLK1. Given these findings, and the possibility that compounds often behave differently in different cellular contexts and may be annotated based on a particular one, it is not surprising that compounds targeting particular genes do not show a consistent directionality relationship with ORFs or CRISPRs.

## Discussion and Limitations

Biology would benefit greatly from the machine learning community turning its attention to rich, single-cell imaging data. While our results may only touch upon the potential applications of machine learning, our emphasis is a strategic appeal to the ML community. We hope that the Resource and benchmarks we created will provide a foundation on which researchers can develop and test novel methods for representation learning, multi-view learning, information retrieval, and style transfer, among others. The task of identifying genes targeted by a given compound to understand its mechanism is exceptionally difficult, expensive, and time-consuming, creating a major bottleneck in developing useful compounds for chemical biology and drug discovery ^11^. This has several implications affecting this work: first, even with low but non-zero success rates, biologists can use their understanding of the compound’s known traits to create and test hypotheses about a chemical’s mechanism of action, accelerating discovery. As well, the predictions from this method might be combined with other predictive approaches, such as rank ordered candidate gene lists from structure based chemical-protein binding predictions, to produce more reliable results. We also emphasize that, as a direct result of the fact that identifying gene targets of compounds is so difficult, only a very limited amount of rather noisy ground truth exists; each known compound-gene interaction was painstakingly discovered after hundreds of thousands of dollars of effort over many years, and many pairings are uncertain. By contrast, many mainstream machine learning (ML) tasks are oriented to replicate specific human skills where ground truth can be collected at large scale (e.g., translation or image recognition), given sufficient resources, and accuracy often approaches 100%. We believe that supervised methods hold promise ^1^, and we hope that novel ML methods developed using our dataset will be used to discover new gene-compound connections that can add to ground truth for this problem in the future. Even with the few hundred perturbations in this dataset, our recent research finds that a transformer-based classification model outperforms the baseline benchmark in this study (unpublished data).

Beyond biology, our dataset provides a challenging, real world test bed for many kinds of more general ML algorithms. It is a large-scale perturbation experiment with complex multidimensional, hierarchical data (images displaying dozens of cells each), and we believe new ML strategies still need to be developed to realize its full potential. In addition to the prediction problems we present in this paper, it also opens up problems in high-level reasoning on experimental data, allowing the study of complex artificial intelligence strategies, such as causal inference (observations from interventional experiments), planning (optimizing the next intervention that maximizes discovery), and simulations (what would have happened if other interventions are applied).

Finally, unusual aspects of the data type that we present pose challenges to ML algorithms and will require they be pushed in different directions to adapt. This may spark creative solutions with broader impact in ML. For example, multiplexed imaging will push the field of machine learning to adapt to domains outside natural images’ red green blue (RGB) colorspace, where the number and relationship among the channels (e.g., the extent of correlation) is very different compared to natural images of everyday objects.

In general, deep learning-based features may provide improvements in performance for some tasks over the classical CellProfiler-based features, though interpretability remains a challenge. The direct interpretability of CellProfiler-based features can help in examining signatures and comparing them to evidence found in existing research relevant to a particular profiling task. One can readily visualize any target cell population by looking up the associated sample cell images or by converting CellProfiler representations into images with CNN-based image generators. An example set of cell images generated by a fundamental version of such a model are shown in Supplementary Figure 2. The research community can delve deeper into creating interpretative models from CellProfiler profiles that we have provided in this resource.

Still, this dataset has limitations. It covers only 160 genes and 303 compounds, and ∼21% of compounds in this dataset are annotated as targeting more than one gene family (Supplementary Figures 3 and 4). Ideally, such a dataset would include only compounds that are very well-studied as targeting one and only one gene’s product. In reality, this is impractical. Polypharmacology is increasingly recognized as common for compounds ^12,13^, and this likely substantially impacts the ability of compounds’ images to match one of the annotated genes’ images. As well, the gene annotations of the compounds may not be complete because of undiscovered gene-compound interactions.

We also note limitations in the selection of genes and compounds. This dataset was curated with compounds and genes available in the Broad’s drug repurposing hub and its library of genetic perturbation reagents, respectively, which introduces several biases. First, the set contains only preclinical/clinical compounds with stronger binding and higher specificity than randomly synthesized compounds; however, for the tasks of mechanism of action determination and virtual screening (where the goal is ultimately to identify such compounds), this is not an overly concerning bias. Second, because of our selection criterion that at least two compounds should target every gene, all compound-gene pairs without a second compound in the repurposing hub were excluded, making better-studied compounds more represented. There were also other selection criteria that introduce various biases, such as, the selected compounds should not be a controlled substance. In terms of experimental conditions, we used a single concentration (5 uM) for all compounds, which is not ideal and, we created this dataset in a single experiment at a single facility; this choice minimizes technical variability and therefore maximizes the biological signal in the data, but also limits the potential for generalizability of any models using it as training data. We note that generalizability of models across datasets is often unnecessary in biology experiments where controls can be included within each experiment; in fact, we recommend those creating large datasets to include the sets of compounds and genes we present here in the experiment to have internal controls/landmarks for the assay. Our consortium has adopted this approach in creating our large-scale dataset of nearly 140,000 chemical and genetic perturbations ^14^.

## Online Methods

### Compound and gene selection

The CPJUMP1 dataset consists of images and profiles of cells that were perturbed separately by chemical and genetic perturbations, where both sets were chosen based on expected “matching” relationships among them. Chemical perturbations are small molecules (compounds) that affect the function of cells while the genetic perturbations are either *open reading frames* (ORFs) that overexpress genes (i.e., yield more of the gene’s product in the cell) or *guide RNAs* that mediate CRISPR-Cas9 (clustered regularly interspaced short palindromic repeats), which knockout gene function (by decreasing production of the gene’s product in the cell – the term knockout is used in the field, although in the timescale of these experiments residual protein likely remains, depending on natural rates of protein turnover).

We therefore designed CPJUMP1 such that for each gene, we have one ORF, two CRISPR guides (for all but 15 genes), and one or two compounds that are thought to impact the cell by influencing the function of that gene’s product (although they may also influence the function of other genes due to polypharmacology, complicating the signal, see Supplementary Figure 4).

We derived the list of compounds from Broad’s Drug Repurposing Hub dataset ^8^, a curated and annotated collection of FDA-approved drugs, clinical trial drugs, and pre-clinical tool compounds (Supplementary Figure 5d). The genes perturbed by genetic perturbations were chosen because they are the annotated targets of the compounds. The specific criteria for compounds, genetic reagents (considering their on- and off-target effects), and controls are described in the section *Compound selection criteria* and their layout on the plates, are described in the section *Plate layout design*. After applying the filters and including controls, we selected a total of 303 compounds and 160 genes such that their corresponding perturbations could fit into three 384-well plates with controls.

### Compound selection criteria

We filtered the Repurposing Hub compounds using several criteria, of which three are most important:

1. The compounds should target genes that belong to diverse gene families (Supplementary Table 7). This is because methods for representation learning and gene-compound matching should work well for many different biological pathways, not just a few that are well-characterized and/or easy to predict.
2. Each gene should be targeted by at least two compounds, so that gene-compound matching and compound-compound matching can both be performed using the dataset.
3. We additionally considered applying the constraint that each compound should target only a single gene. However, this criterion is difficult to achieve due to polypharmacology (Supplementary Figure 4), which is the property for compounds to bind and impact many different gene products in the cell; this is especially common for protein kinase inhibitors in the dataset. Instead, we only filtered out the so-called “historical compounds” listed in the Chemical Probes Portal ^15^, which are compounds that are known to be quite non-selective (or not sufficiently potent) compared with other available chemical probes.

Our list of compounds and genes also includes both negative and positive controls. The negative controls for each perturbation modality are:

● Compounds: DMSO (Dimethyl sulfoxide), which is the solvent for all the compounds studied. In other words, all samples will have DMSO added at the same concentration, but the negative controls have no *additional* compound added.
● ORFs: 15 ORFs with the weakest signature in previous image-based profiling experiments ^16^. Thus, the total number of genes with ORFs is 160+15=175.
● CRISPR guides: 30 CRISPR guides that target an intergenic site (cutting controls, n = 3) or don’t have a target sequence that exists in human cells (non-cutting controls, n = 27).

There are three types of compound positive controls in our list. First, we included chemical probes that are very well-studied and (unlike most compounds) are known to very selectively modulate the genes that they target (poscon_cp) ^15^. Second, we included compounds that strongly correlate with the correct genetic perturbation in previous image-based profiling experiments with ORFs ^16^ and compounds (poscon_orf) ^17^. Finally, we included a set of very diverse pairs of compounds with strong intra-pair and weak inter-pair correlations, based on prior experiments (poscon_diverse).

Additionally, compounds were filtered based on availability through at least one of four compound vendors (Sigma, SelleckChem, Tocris, and MedChemEx) and genetic reagents through Broad’s Genetic Perturbation Platform portal. Lastly, we also excluded compounds on the U.S. Drug Enforcement Agency (DEA) list of controlled substances or the Organisation for the Prohibition of Chemical Weapons (OPCW) list of chemical weapons precursors.

### Target loci selection

We picked the target loci for the CRISPR experiments by selecting the top-two-ranking sgRNA sequences that maximize their on-target activity, calculated using the Azimuth 2.0 model ^18^ and minimize the off-target activity, calculated using the Cutting Frequency Determination score (additional details can be found at https://portals.broadinstitute.org/gpp/public/software/sgrna-scoring-help).

### Compound and gene metadata

A list of CRISPR reagents and their target sequences is available here: https://github.com/jump-cellpainting/2023_Chandrasekaran_submitted/blob/5c6fcf9dc70a85176f5afc5263acbc230d90ca40/metadata/external_metadata/JUMP-Target-1_crispr_metadata.tsv

A list of ORF reagents and their target sequences is available here:

https://github.com/jump-cellpainting/2023_Chandrasekaran_submitted/blob/5c6fcf9dc70a85176f5afc5263acbc230d90ca40/metadata/external_metadata/JUMP-Target-1_orf_metadata_with_sequence.tsv

A list of compounds with their names, PubChem ID, SMILES and gene targets is available here:

https://github.com/jump-cellpainting/2023_Chandrasekaran_submitted/blob/5c6fcf9dc70a85176f5afc5263acbc230d90ca40/metadata/external_metadata/JUMP-Target-1_compound_metadata_targets.tsv

Infection efficiency data for the ORF and CRISPR regents for each time point and cell type from CellTiter-Glo cell viability assay is available here:

https://github.com/jump-cellpainting/2023_Chandrasekaran_submitted/blob/5c6fcf9dc70a85176f5afc5263acbc230d90ca40/metadata/external_metadata/CPJUMP1_Infection_Efficiency.xlsx

### Plate layout design

The first plate (Supplementary Figure 5a) is entirely **compounds**, with two compounds per gene target. Each compound is in singlicate on the plate except for a dozen or so compounds (poscon_diverse) in duplicate and the negative control DMSO described above, in n=64 replicates. The second plate (Supplementary Figure 5b) is entirely **CRISPR** reagents, with two guides per gene, each arrayed in its own well and kept separate, with no within-plate replicates; there are two replicates of the 30 CRISPR negative controls described above. The third plate (Supplementary Figure 5c) is entirely **ORFs**; because there was only one perturbation reagent per gene, there are two replicates of each per plate, plus n=4 replicates of the 15 ORF negative controls. Each plate contains only one type of perturbation modality.

We also considered the impact of edge effects, or plate-layout effects, in our design. Edge effects are the technical artifact whereby different samples will yield different behavior depending on where they are located on a plate; generally this is most observed in the outer two rows and columns of the plate, and the problem persists despite efforts to mitigate it experimentally ^19^. While designing the plate layout, we divided the plate into outer and inner wells, where the outer wells are the two rows and columns closest to the edge of the plate and the inner wells are the rest of the wells on the plate. Then we applied the following constraints to minimize the impact of edge effects:

● Both of the compounds that target the same gene will either be in the inner wells or in the outer wells. They will not be split such that one of the compounds is in the inner well while the other is in the outer well.
● The gene target of outer well compounds will be in the outer wells of the genetic perturbation plate.
● All the positive control compounds are in the inner wells.

If preferable, with this design, an analysis can be constrained to the inner wells only, to ensure that edge effects have minimal influence on the results.

### Experimental conditions Compound

● Concentration: The treatment compounds were assayed at 5 uM
● Cell seeding density: 1000 cells/well

### ORF

● Cell seeding density: 1625 cells/well, 40 uL media seeding volume/384w
● Viral volume: 1 uL virus/384w
● Polybrene: 4 ug/mL
● Media change after 24 hrs removing polybrene and virus adding back 50 uL media
● No selection with Blasticidin.

### CRISPR

● Cell seeding density: 350 cells/well, 40 uL media seeding volume/384w,
● Viral volume: 0.5 uL virus/384w
● Polybrene: 4 ug/mL polybrene
● Media change after 24 hrs removing polybrene and virus adding back 50 uL media
● No selection with Blasticidin.

### Experimental conditions tested

Although constrained by cost, we captured the compound, ORF, and CRISPR plates under several experimental conditions to identify those that improve gene-compound, compound-compound and gene-gene matching (Figure 3). We did more replicate plates for conditions that were less expensive or that were the most promising, and for the compound and CRISPR plates which had only singlicates of most samples (as compared to ORFs which had duplicates within the plate). Additionally, we captured plates under many other experimental conditions, listed below, for optimizing the experimental conditions. UMAP embeddings of all the experimental conditions are shown in Supplementary Figures 6-11.

We note that the cell types are commonly used historical lines derived from two white patients, one male (A549) and one female (U2OS). Therefore, conclusions from this data may only hold true for the demographics or genomics of those persons and not broader groups. They were chosen because the lines are both well-suited for microscopy, and they offer the advantage of enabling direct comparison to extensive prior studies using them.

### Full list of experimental conditions

Primary group of experimental conditions

1. Four replicate plates of compounds and CRISPR guides and two replicate plates of ORFs (which, as mentioned, contain two replicates within each plate) at two time points and two cell lines each. The short and long time points were different for each perturbation type: compounds (24-hour, 48-hour), ORFs (48-hour, 96-hour) and CRISPR guides (96-hour, 144-hour). The two cell lines were U2OS and A549. For CRISPR experiments, polyclonal A549 and U2OS were used.

Secondary group of experimental conditions

2. One plate of the A549 96-hour ORF plate, where the cells have been additionally treated with Blasticidin (a drug that kills cells that have not been properly infected with the genetic reagent).
3. Two replicate plates of the A549 144-hour CRISPR plate, where the cells have been additionally treated with Puromycin (a drug that kills cells that have not been properly infected with the genetic reagent).
4. Two replicate plates of the A549 48-hour compound plate with 20% higher cell seeding density than the baseline.
5. Two replicate plates of the A549 48-hour compound plate with 20% lower cell seeding density than the baseline.
6. Four replicate plates of the A549 24-hour compound plate were imaged six additional times to test photobleaching from repeated imaging.
7. Two replicates of the ORF plates in U2OS and A549 at 96-hour and 144-hour were imaged four additional times, once on each of days 1, 4, 14, 28 after the first imaging, to test the stability of samples over time.
8. Four replicate plates of 48-hour compound plates in polyclonal A549 cells with Cas9.

### Number of plates and images

Across both the groups of experimental conditions, there are 51 physical plates and 107 plates of images, 40 physical plates in the primary group and 11 physical plates and 67 plates of images in the secondary group. Each plate consists of 384 wells and on average, nine sites were imaged within each well. At each site, eight (five fluorescent and three brightfield) images were captured. This amounts to nearly three million images across the 107 plates.

### Sample preparation and image acquisition

The Cell Painting assay involves staining eight components of cells with six fluorescent dyes that are imaged in five channels: nucleus (Hoechst; DNA), nucleoli and cytoplasmic RNA (SYTO 14; RNA), endoplasmic reticulum (concanavalin A; ER), Golgi and plasma membrane (wheat germ agglutinin (WGA); AGP), mitochondria (MitoTracker; Mito), and the actin cytoskeleton (phalloidin; AGP) (Figure 1). We optimized the Cell Painting assay described in Bray et al. (2016) by changing the concentrations of Hoechst, phalloidin, and combining dye addition and dye permeabilization steps to create Cell Painting v2.5. The changes to the protocol are listed at https://github.com/carpenterlab/2022_Cimini_NatureProtocols/wiki#changes-in-the-official-protocol-to-create-v25-chandrasekaran-et-al-2021. The changes are described in more detail in Cimini et al. (2022), in which we further developed Cell Painting v3, where we also changed the concentrations of concanavalin A and SYTO14. The images were acquired across five fluorescent channels plus three brightfield planes using a Revvity Opera Phenix HCI microscope in widefield mode with 16-bit depth and with a 20x/1.0NA water immersion lens. 2x2 pixel binning was used, for a final effective pixel size of 0.598 µm.

### Image display

In Figure 1, each channel was mapped to a final 0-255 LUT per the following colors and display ranges in Fiji ^20^ – Ch1 (Mito): Red, (1078-10191); Ch2 (AGP): Orange Hot, (426-22225); Ch3 (RNA): Yellow, (360-33716); Ch4 (ER): Green, (238-14272); Ch5 (DNA): Cyan, (238-20508). The final image represents the max value in the RGB color-space for all 5 channels, created with Fiji’s “Composite (Max)” mode. No other (linear or nonlinear) adjustments were performed. Panels were assembled using the “Magic Montage” Fiji tool and channel annotations added in Google Draw.

### Image processing

We used CellProfiler bioimage analysis software (version 4.0.6) to process the images using classical algorithms ^21^. We corrected for variations in background intensity^22^, and then segmented cells, distinguishing between nuclei and cytoplasm. Then, across the various channels captured, we measured various features of cells across several categories including fluorescence intensity, texture, granularity, density, location (see http://cellprofiler-manual.s3.amazonaws.com/CellProfiler-4.0.6/index.html for more details). Following the image analysis pipeline (see https://github.com/jump-cellpainting/2023_Chandrasekaran_submitted/tree/5c6fcf9dc70a85176f5afc5263acbc230d90ca40/pipelines for the pipelines), we obtained 5792 feature measurements from each of more than 75 million cells.

### Image-based profiling

We used cytominer and pycytominer workflows to process the single cell features extracted using CellProfiler ^23–26^. We aggregated the single cell profiles by computing the median profile, and then normalized the averaged profiles by subtracting the median and dividing by the median absolute deviation (m.a.d.) of each feature. This was done in two ways: using the median and m.a.d. of (i) the negative control wells on the plate (used in the analysis shown here), and (ii) all the wells on the plate. Finally, we filtered out redundant features (such that no pair of features has Pearson correlation greater than 0.9) as well as features with near-zero variance across all the plates.

### Average Precision (AP), mean Average Precision (mAP) and Fraction Retrieved (FR)

*Average precision* (AP) serves as our primary metric for both replicability (how distinguishable are replicates of a perturbation from negative controls) and biological relevance (how distinguishable are sister perturbations from other perturbations). We measure similarity between profiles using cosine similarity or absolute value of cosine similarity for cases where both positive and negative correlations are considered matches. To calculate AP, we first construct a binary ranking of sample and negative control profiles by their cosine similarities. Based on this rank list, calculation of the AP score follows the common formulation in the field ^27^, i.e. we average precision values at each rank *k* where recall changes:

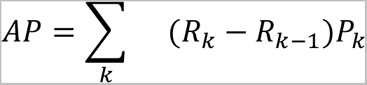

We then assign a *p*-value to each AP score by a permutation-based significance testing approach. Specifically, we assess the significance of the AP score against a null distribution built by randomly shuffling the rank list 100,000 times and computing corresponding random AP scores. These *p*-values are then adjusted for multiple testing using the Benjamini-Hochberg procedure to obtain *q*-values.

We also compute *mean Average Precision* (mAP) by averaging AP scores per each class, where “class” refers to either a specific perturbation (for replicability) or a gene targeted by compound or genetic perturbation (for biological relevance). We report a per-class mAP value along with a combined *q*-value obtained by taking the geometric mean of the q-values. Finally, we summarize the mAP values for a specific task by calculating the fraction of classes with *q*-values below the significance threshold (0.05), termed *Fraction Retrieved* (FR).

### Recommended dataset splits

The methods presented in the benchmarks do not involve any training (we simply use a predetermined similarity metric and hand-engineered features or a pre-trained model) and thus did not require creating the typical train-validate-test data splits. Depending on the use case, we suggest using different splits. For the two benchmarks, we offer the following guidelines for creating data splits when training is involved:

### Representation learning

For a general representation learning and domain adaptation tasks, one could train on the dataset from one cell line or time point and test it on the other cell line or time point.

### Gene-compound matching

1. All replicates of a perturbation should be in the same split.
2. For the CRISPR dataset, both guides should be in the same split.
3. Three of the compounds (BVT-948, dexamethasone, and thiostrepton) have two different identifiers each in the dataset (because of small differences in structures) but the same compound name. Each pair should be in the same split.
4. If analyzing data at the single cell level, all cells from a well should be in the same split.

We provide recommended data splits in https://github.com/jumpcellpainting/2023_Chandrasekaran_submitted/tree/main/datasplits

### Code and Data availability

Well-level morphological profiles, image analysis pipelines, profile generation pipelines, plate maps and plate and compound metadata, and instructions for retrieving the cell images and single cell profiles are publicly available online at https://broad.io/cpjump1. The code for reproducing the benchmark results, tables, and figures are available at https://github.com/jump-cellpainting/2023_Chandrasekaran_submitted/tree/5c6fcf9dc70a85176f5afc5263acbc230d90ca40/benchmark and https://github.com/jump-cellpainting/2023_Chandrasekaran_submitted/tree/5c6fcf9dc70a85176f5afc5263acbc230d90ca40/visualization.

The landing page of the GitHub repository for this dataset has relevant additional information: https://broad.io/cpjump1.

Cell Painting images and single-cell profiles are available at the Cell Painting Gallery on the Registry of Open Data on AWS (https://registry.opendata.aws/cellpainting-gallery/) under accession number cpg0000-jump-pilot. For well-level aggregated profiles, we use GitHub as the hosting platform and the files are stored in GitLFS.

We have released the data with a CC0 license and the code with a BSD 3-Clause license.

### Tools and Software used

Data analysis was performed using python code written in a Jupyter notebook environment ^28,29^. Python libraries used for data analysis include Numpy, Scipy, scikit-learn and pandas ^30–33^. Plots were generated using matplotlib, seaborn and Plotly libraries ^34–36^. Fiji^20^’s Magic Montage plugin, Lucidchart and Google Draw were used for generating montages, creating schematics and for adding text to images. Two-dimensional representations of image-based profiles in Figures 2 and Supplementary figures 5-11 were generated using UMAP^37^ .

## Acknowledgements and Disclosure of Funding

The authors appreciate the more than 100 scientists who have contributed to the organization and scientific direction of the JUMP Cell Painting Consortium. The authors acknowledge valuable inputs on data consistency from Srijit Seal and David Figueiredo Vidal.

The authors gratefully acknowledge a grant from the Massachusetts Life Sciences Center Bits to Bytes Capital Call program for funding the data production. We appreciate funding to support data analysis and interpretation from members of the JUMP Cell Painting Consortium, from the National Institutes of Health (NIH MIRA R35 GM122547 to AEC), and from the Chan Zuckerberg Initiative DAF, an advised fund of the Silicon Valley Community Foundation (2020-225720 to BAC). The authors also gratefully acknowledge the use of the Revvity Opera Phenix High-Content/High-Throughput imaging system at the Broad Institute, funded by the S10 Grant NIH OD-026839-01.

