## Supplementary materials for "Three million images and morphological profiles of cells treated with matched chemical and genetic perturbations"

| **Perturbation modality** | **Time point – short** | **Time point – long** |
| --- | --- | --- |
| Compound | 24-hour | 48-hour |
| ORF | 48-hour | 96-hour |
| CRISPR | 96-hour | 144-hour |

Supplementary Table 1: **Perturbation time points**. Description of the *short* and *long* time points for each perturbation modality.

| **Type** | **Description** | **No. of data points** |
| --- | --- | --- |
| Replicate retrieval | compound_long_A549 | 306 |
| Replicate retrieval | compound_long_U2OS | 306 |
| Replicate retrieval | compound_short_A549 | 306 |
| Replicate retrieval | compound_short_U2OS | 306 |
| Replicate retrieval | crispr_long_A549 | 305 |
| Replicate retrieval | crispr_long_U2OS | 305 |
| Replicate retrieval | crispr_short_A549 | 305 |
| Replicate retrieval | crispr_short_U2OS | 305 |
| Replicate retrieval | orf_long_A549 | 160 |
| Replicate retrieval | orf_long_U2OS | 160 |
| Replicate retrieval | orf_short_A549 | 160 |
| Replicate retrieval | orf_short_U2OS | 160 |
| Sister perturbation retrieval | compound_long_A549 | 402 |
| Sister perturbation retrieval | compound_long_U2OS | 285 |
| Sister perturbation retrieval | compound_short_A549 | 315 |
| Sister perturbation retrieval | compound_short_U2OS | 300 |
| Sister perturbation retrieval | crispr_long_A549 | 73 |
| Sister perturbation retrieval | crispr_long_U2OS | 42 |
| Sister perturbation retrieval | crispr_short_A549 | 73 |
| Sister perturbation retrieval | crispr_short_U2OS | 73 |
| Gene compound retrieval | compound_long_crispr_long_A549 | 134 |
| Gene compound retrieval | compound_long_crispr_long_U2OS | 92 |
| Gene compound retrieval | compound_long_crispr_short_A549 | 135 |
| Gene compound retrieval | compound_long_crispr_short_U2OS | 115 |
| Gene compound retrieval | compound_long_orf_long_A549 | 8 |
| Gene compound retrieval | compound_long_orf_long_U2OS | 29 |
| Gene compound retrieval | compound_long_orf_short_A549 | 8 |
| Gene compound retrieval | compound_long_orf_short_U2OS | 53 |
| Gene compound retrieval | compound_short_crispr_long_A549 | 128 |
| Gene compound retrieval | compound_short_crispr_long_U2OS | 102 |
| Gene compound retrieval | compound_short_crispr_short_A549 | 130 |
| Gene compound retrieval | compound_short_crispr_short_U2OS | 125 |
| Gene compound retrieval | compound_short_orf_long_A549 | 8 |
| Gene compound retrieval | compound_short_orf_long_U2OS | 30 |
| Gene compound retrieval | compound_short_orf_short_A549 | 8 |
| Gene compound retrieval | compound_short_orf_short_U2OS | 58 |

Supplementary Table 2: Number of data points in each box plot in Figure 4.

| **Cell** | **Fisher’s exact statistic** | **p value** |
| --- | --- | --- |
| U2OS | 2.64156 | <0.05 |
| A549 | 2.19389 | <0.05 |

Supplementary Table 3: **Comparing the directionality of ORF-CRISPR cosine similarities.** We performed Fisher’s exact test to determine whether the cosine similarities between ORFs and CRISPR guides were positive or negative. ORF-CRISPR pairs that were beyond the 5th and 95th percentile of the null distribution (cosine similarities of ORF and CRISPR guides targeting different genes) were used in the Fisher’s test. A positive, significant (p <0.05) statistic means that the ORF-CRISPR cosine similarities are more positive than negative.

| **Cell** | **Genetic Perturbation** | **Fisher’s exact statistic** | **p value** |
| --- | --- | --- | --- |
| A549 | CRISPR | 2.03487 | <0.05 |
| A549 | ORF | 2.19404 | <0.05 |
| U2OS | CRISPR | 2.81489 | <0.05 |
| U2OS | ORF | 2.63868 | <0.05 |

Supplementary Table 4: **Comparing the directionality of Compound-CRISPR and Compound-ORF cosine similarities.** We performed Fisher’s test to determine whether the cosine similarities between compounds and each genetic perturbation type (ORF and CRISPR guides) were positive or negative. Compound-genetic perturbation pairs that were beyond the 5th and 95th percentile of the null distribution (cosine similarities of compound-genetic perturbations targeting different genes) were used in the Fisher’s test. A positive, significant (p <0.05) statistic means that the compound-genetic perturbation cosine similarities are more positive than negative.

| **Cell** | **Genetic**  **Perturbation** | **pert_iname** | **Metadata_matching_target** | **moa_list** | **Cosine similarity** |
| --- | --- | --- | --- | --- | --- |
| U2OS | CRISPR | BI-2536 | PLK1 | PLK inhibitor | 0.779615 |
| U2OS | CRISPR | AMG900 | AURKB | Aurora kinase inhibitor | 0.72932 |
| U2OS | CRISPR | danusertib | AURKB | Aurora kinase inhibitor, growth factor receptor inhibitor | 0.654305 |
| U2OS | ORF | PP-2 | ABL1 | src inhibitor | 0.643488 |
| U2OS | ORF | ponatinib | LYN | Bcr-Abl kinase inhibitor, FLT3 inhibitor, PDGFR tyrosine kinase receptor inhibitor | 0.61835 |
| U2OS | ORF | BI-2536 | BRD4 | PLK inhibitor | 0.598121 |
| U2OS | ORF | pentostatin | ADA | adenosine deaminase inhibitor, ribonucleotide reductase inhibitor | 0.583311 |
| U2OS | ORF | hexestrol | AKR1C1 | synthetic estrogen | 0.5711 |
| U2OS | ORF | pyrrolidine-dithiocarbamate | HSD11B1 | NFkB pathway inhibitor | 0.566448 |
| U2OS | CRISPR | sodium-butyrate | FFAR2 | HDAC inhibitor | 0.560779 |
| U2OS | CRISPR | TG-003 | CLK1 | CLK inhibitor | -0.326471 |
| U2OS | CRISPR | GSK3787 | PPARD | PPAR receptor antagonist | -0.338408 |
| U2OS | ORF | amlexanox | FGF1 | histamine receptor modulator | -0.343726 |
| U2OS | ORF | pazopanib | FGF1 | KIT inhibitor, PDGFR tyrosine kinase receptor inhibitor, VEGFR inhibitor | -0.354573 |
| U2OS | CRISPR | PF-06463922 | ALK | ALK tyrosine kinase receptor inhibitor | -0.372329 |
| U2OS | ORF | sotrastaurin | PRKCE | PKC inhibitor | -0.383463 |
| U2OS | CRISPR | benzamil | ASIC1 | sodium channel blocker | -0.424186 |
| U2OS | CRISPR | ZM-336372 | LCK | RAF inhibitor | -0.44488 |
| U2OS | ORF | GSK2110183 | AKT1 | AKT inhibitor | -0.481628 |
| U2OS | ORF | ZM-336372 | MAPK14 | RAF inhibitor | -0.547655 |

Supplementary Table 5: **Compound-genetic perturbation similarity in U2OS cells**. The top 10 positively correlated and negatively correlated compound-genetic perturbations pairs in U2OS. pert_iname is the name of the compound, Metadata_matching_target refers to the target of the compound and genetic perturbation and moa_list is the list of mechanism of action annotation(s) for each compound.

| **Cell** | **Genetic**  **Perturbation** | **pert_iname** | **Metadata_matching_target** | **moa_list** | **Cosine similarity** |
| --- | --- | --- | --- | --- | --- |
| A549 | CRISPR | BI-2536 | PLK1 | PLK inhibitor | 0.757815 |
| A549 | CRISPR | AMG900 | AURKB | Aurora kinase inhibitor | 0.729485 |
| A549 | CRISPR | NSC-663284 | CDC25A | CDC inhibitor | 0.720625 |
| A549 | CRISPR | KI-16425 | LPAR1 | lysophosphatidic acid receptor antagonist | 0.680091 |
| A549 | CRISPR | danusertib | AURKB | Aurora kinase inhibitor, growth factor receptor inhibitor | 0.621753 |
| A549 | CRISPR | fludarabine-phosphate | DCK | ribonucleotide reductase inhibitor | 0.618721 |
| A549 | CRISPR | GSK1070916 | AURKB | Aurora kinase inhibitor | 0.58189 |
| A549 | CRISPR | bepridil | TNNC1 | calcium channel blocker | 0.557164 |
| A549 | CRISPR | carzenide | CA14 |  | 0.55219 |
| A549 | CRISPR | aminopurvalanol-a | CDK2 | CDK inhibitor, tyrosine kinase inhibitor | 0.534492 |
| A549 | ORF | pazopanib | FGF1 | KIT inhibitor, PDGFR tyrosine kinase receptor inhibitor, VEGFR inhibitor | -0.35889 |
| A549 | CRISPR | citric-acid | AKR1B1 | coagulation factor inhibitor | -0.36625 |
| A549 | ORF | ibutilide | KCNH7 | potassium channel blocker | -0.369209 |
| A549 | CRISPR | JTE-607 | TNF | cytokine production inhibitor | -0.377579 |
| A549 | CRISPR | salicylic-acid | AKR1C1 | cyclooxygenase inhibitor | -0.391766 |
| A549 | CRISPR | epoprostenol | PTGIS | prostacyclin analog | -0.395514 |
| A549 | CRISPR | rifamycin | SLCO2B1 | DNA directed RNA polymerase inhibitor | -0.405275 |
| A549 | ORF | sulfasalazine | SLC7A11 | cyclooxygenase inhibitor | -0.417641 |
| A549 | CRISPR | ibutilide | CACNG1 | potassium channel blocker | -0.492405 |
| A549 | CRISPR | sorbinil | AKR1B1 | aldose reductase inhibitor | -0.541175 |

Supplementary Table 6: **Compound-genetic perturbation similarity in A549 cells**. The top 10 positively correlated and negatively correlated compound-genetic perturbations pairs in A549. pert_iname is the name of the compound, Metadata_matching_target refers to the target of the compound and genetic perturbation and moa_list is the list of mechanism of action annotation(s) for each compound.

| Number of gene targets (N) | Number of gene families with N gene targets in the final list |
| --- | --- |
| 1 | 92 |
| 2 | 16 |
| 3 | 2 |

Supplementary Table 7: **Number of gene families with a given number of gene targets.** Closely related genes are called *families.* Gene family names were downloaded from <https://www.genenames.org/download/custom/>. To maximize the diversity of genes and reduce the frequency of retrieving a gene’s family member as an “incorrect” match for a given compound, the 130 genes chosen for the experiment belonged to 110 diverse gene families. The additional 30 genes in the experiment were targets of the positive control compounds in the experiment.

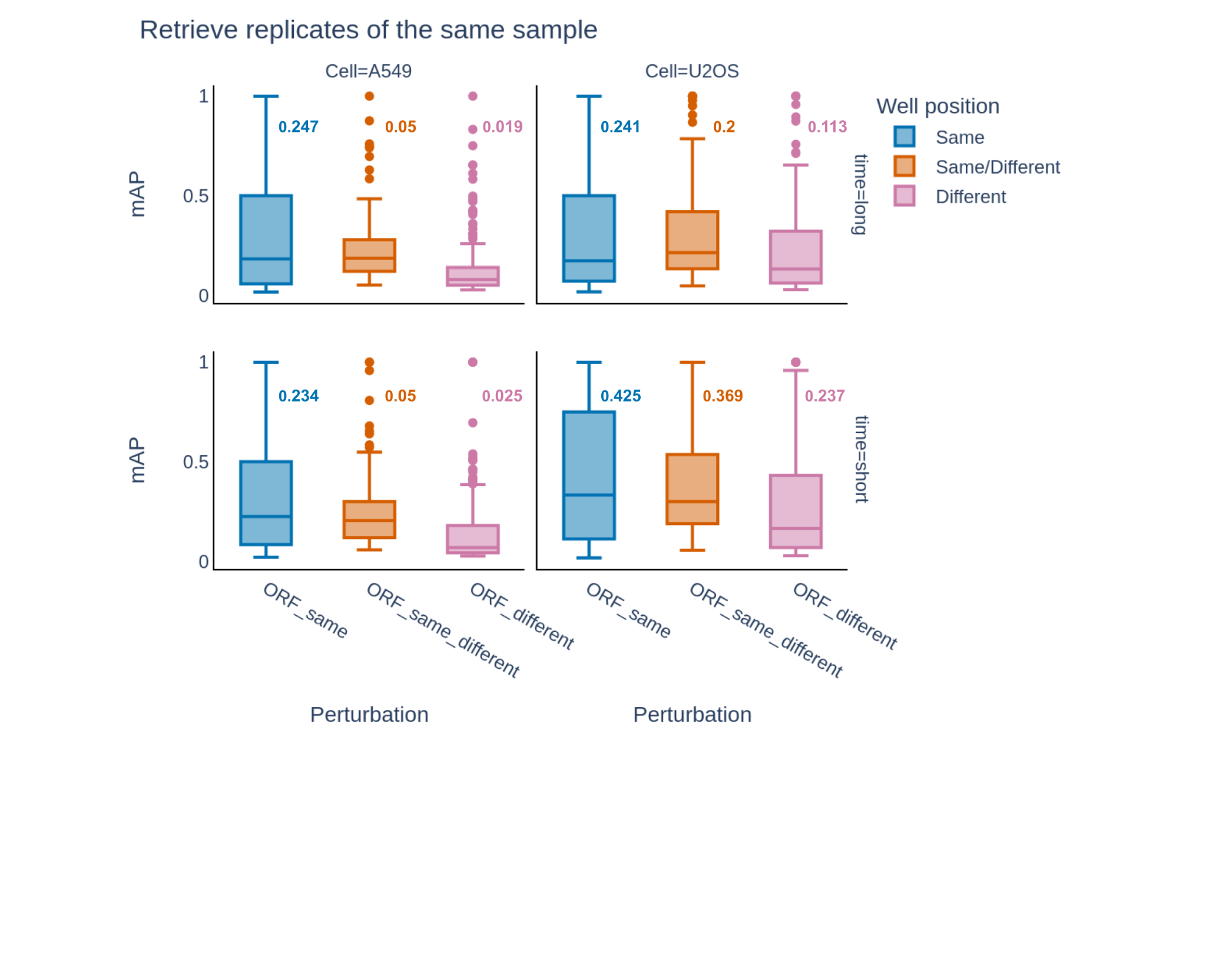

Supplementary Figure 1: **Well position effect.** mAP for perturbation detection for ORFs in the same well position (blue), same or different well positions (red) and different well positions (pink); the same or different well positions is what is shown in Figure 4a in the main text. ORFs in different well positions are affected by plate layout effects, which lowers mAP and FR scores for retrieving replicates against a background of negative control wells.

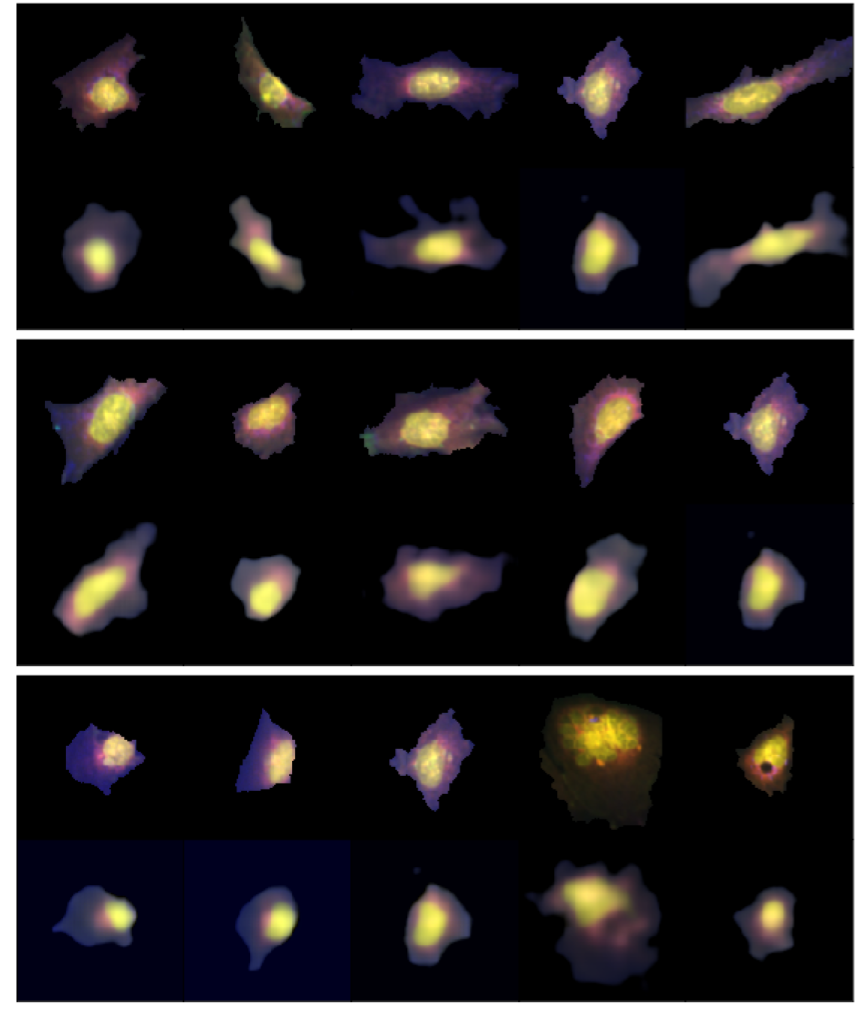

Supplementary Figure 2: **CellProfiler features to cell images.** Example single cell images (first row) and their synthetically generated version (second row) are shown in each sub figure.

The synthetic version is generated by each single cell’s corresponding CellProfiler measurements. To learn a transformation function from single-cell’s CellProfiler extracted features to single cell images, 7077 single cells were randomly selected from a set of eight diverse compounds (aloxistatin, AMG900, dexamethasone, TC-S-7004, FK-866, LY2109761, NVS-PAK1-1 and quinidine) to train a Convolutional Neural Network (CNN). The set of cell-level Cell Painting measurements were reduced to a non-redundant set of features for five channels of *DNA, RNA, ER, AGP* and *Mito*. Location related features and low variance features were excluded. Single-cell images corresponding to each cell’s CellProfiler measurements were extracted by image crops of a fixed size (160 pixels) bounding box around the cell's *Cells_Location_Center* coordinates. CNN model learns the transformation from (3019,1) size CellProfiler features to (128, 128, 5) size images.

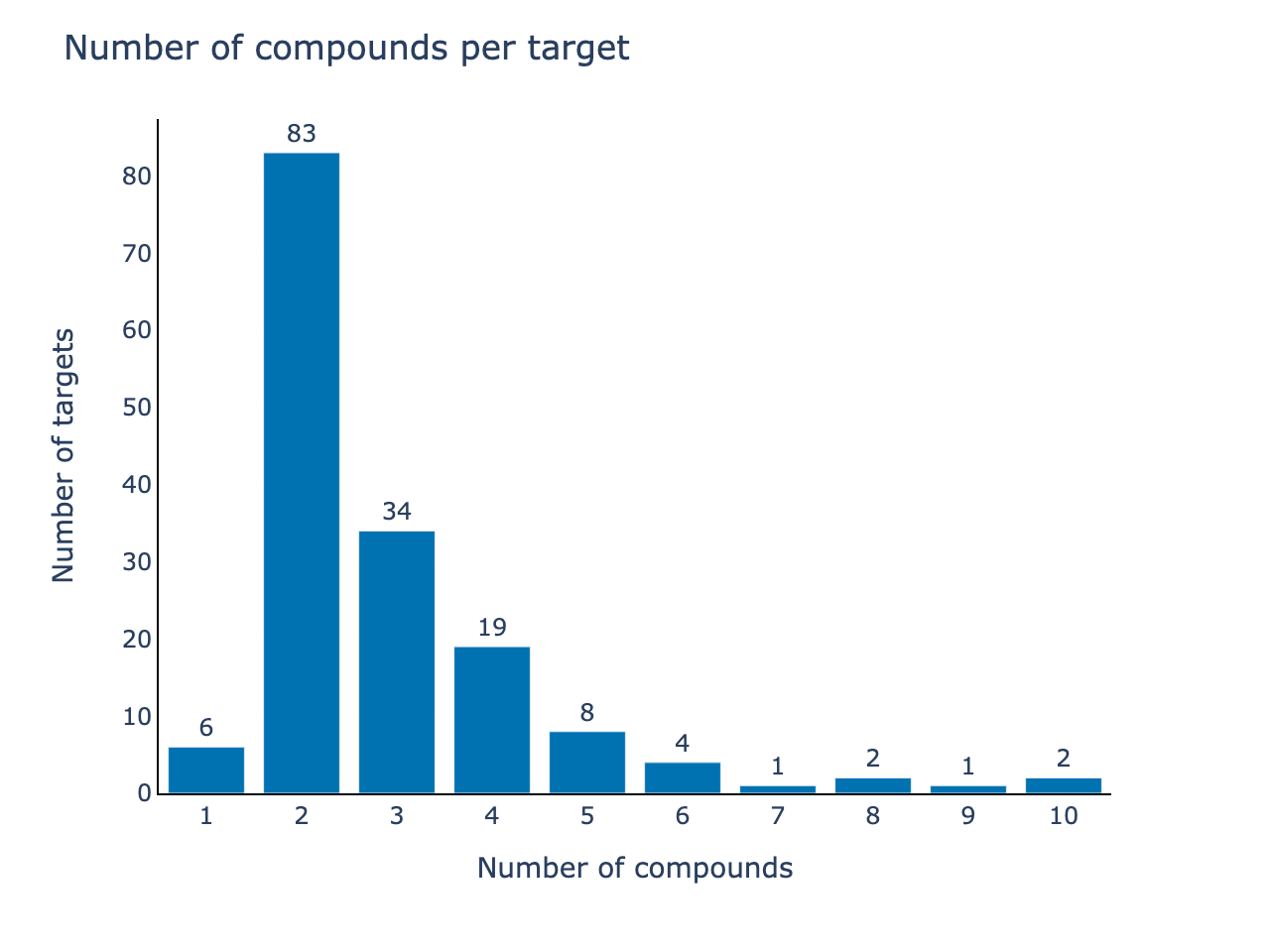

Supplementary Figure 3: **Number of genes targeted by a given number of compounds**. Most genes are targeted by two compounds, though some genes are targeted by as many as ten compounds because many compounds in the set are annotated as having multiple gene targets (see Supplementary Figure 4).

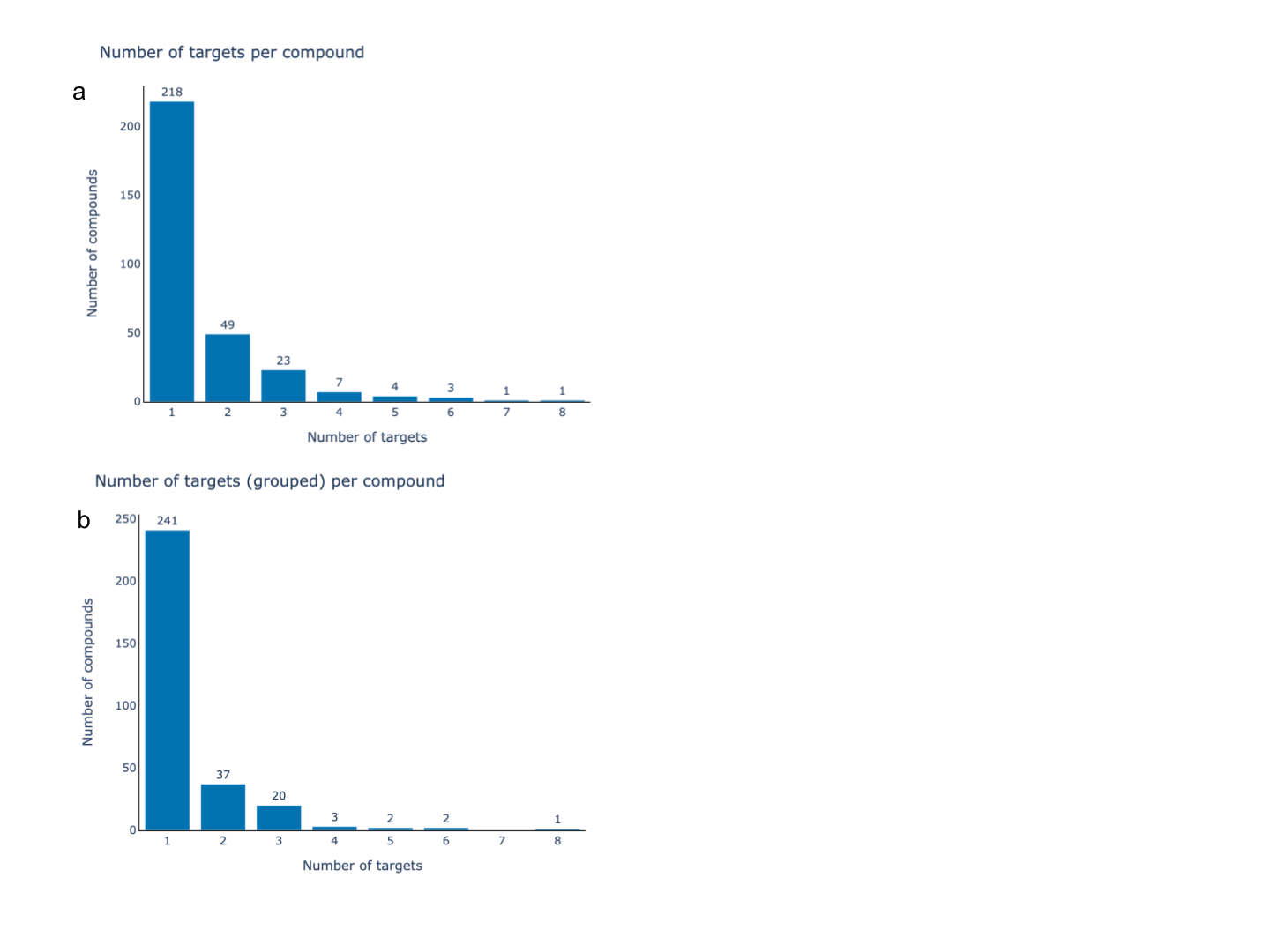

Supplementary Figure 4: **Polypharmacology**. a) Most compounds (n=218) are annotated as targeting a single gene, but there are sixteen compounds that target four or more genes. Because annotations are incomplete, there may be more targets per compound than noted here. b) When members of the same gene family are grouped together for this analysis, for example NTRK1, NTRK2 and NTRK3, then only eight compounds target four or more genes or gene families; the majority of compounds (241) target only a single gene or gene family.

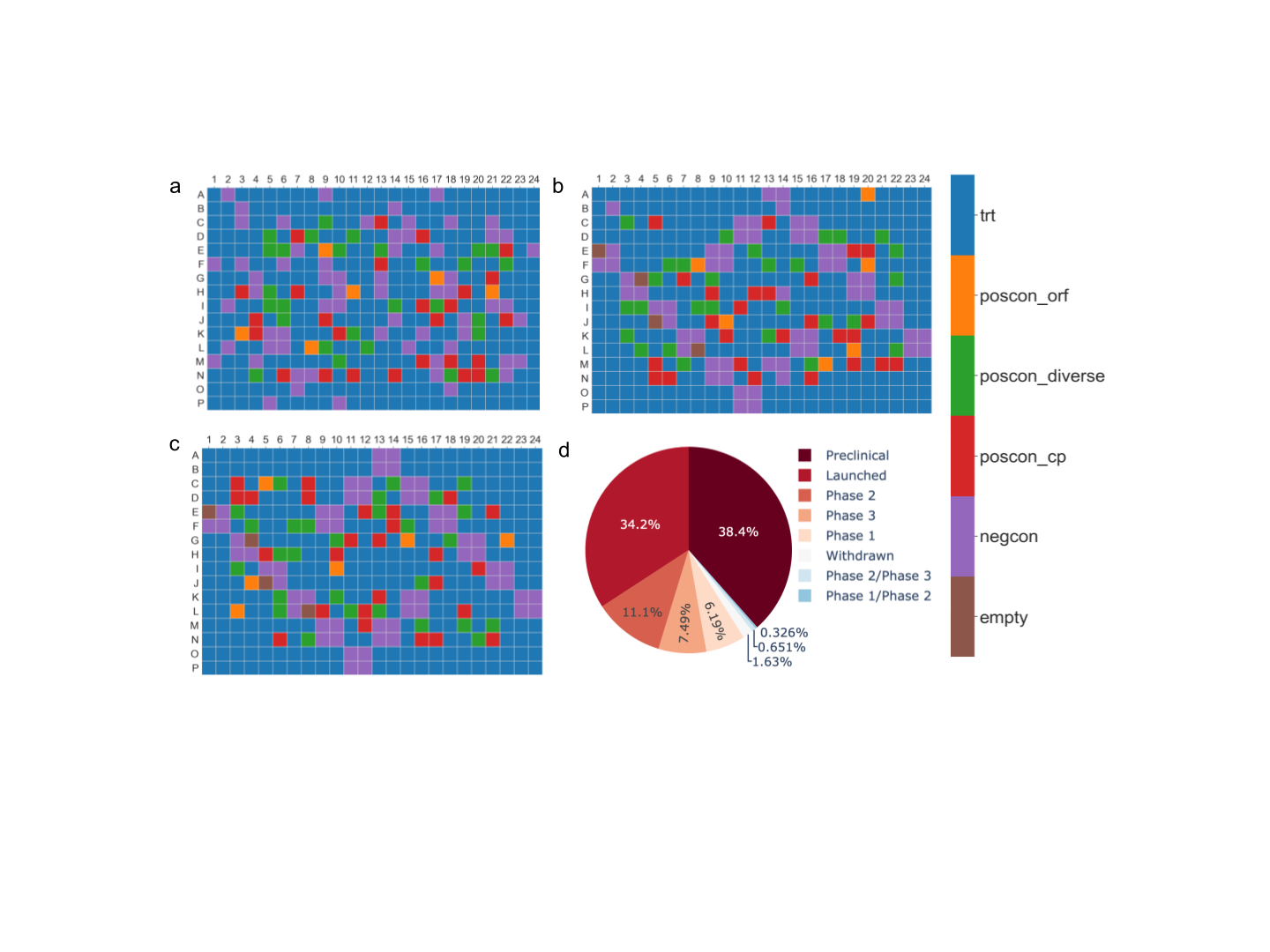

Supplementary Figure 5: **Plate maps and overview of compounds’ clinical phase status**. Maps in a-c show a) Compound plate, b) CRISPR plate and c) ORF plate. The control wells and the treatment (trt) wells are shown in different colors. Poscon are positive controls (additional details in the online methods) and negcon is the negative control. d) Over a third of the compounds in the dataset have been launched for sale, whereas others have progressed to various stages of human clinical trials.

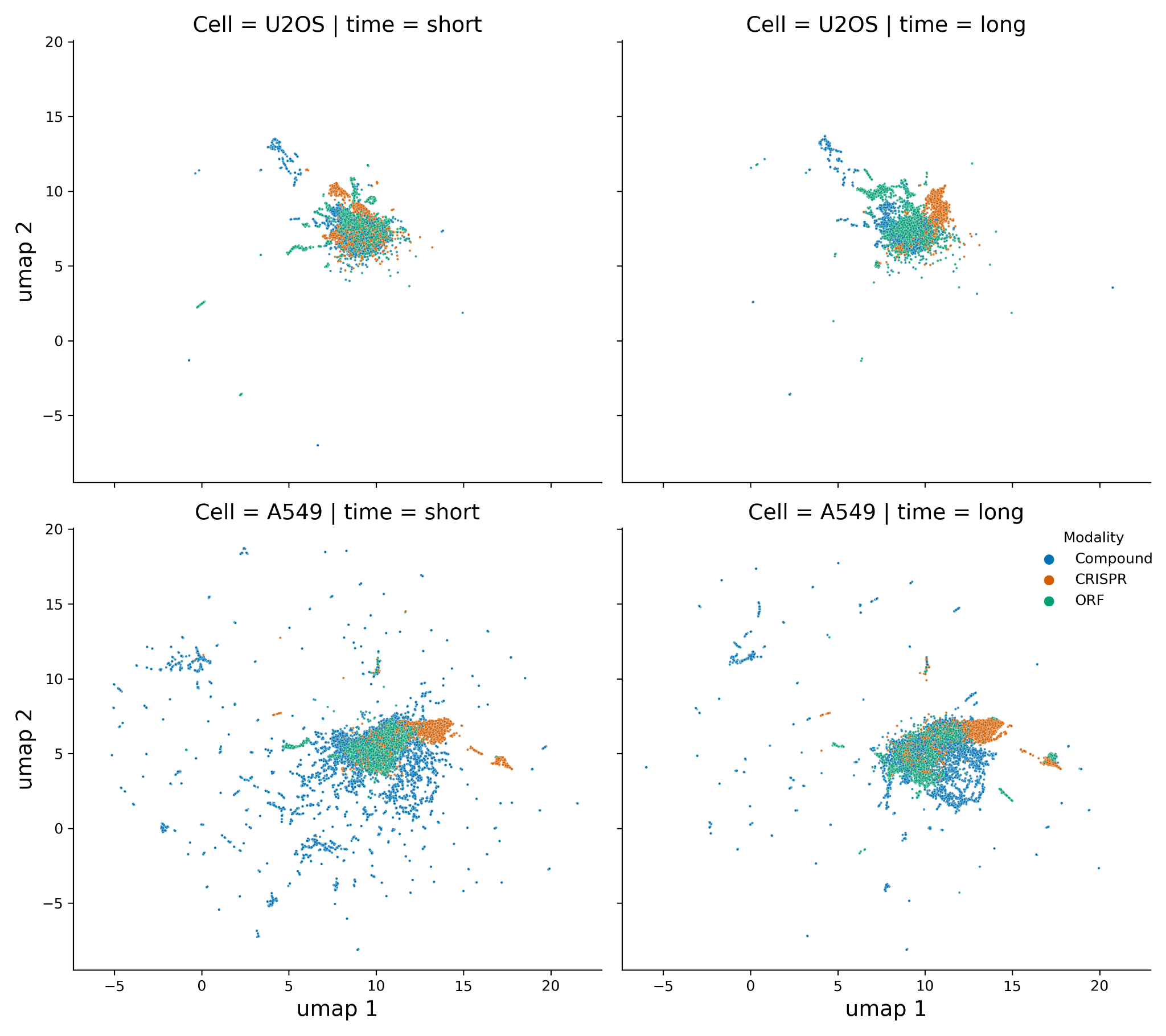

Supplementary Figure 6: **Cell type and time, for all tested perturbations and conditions**. The primary group of tested samples in the CPJUMP1 experiment consists of three perturbation modalities (compounds, CRISPR guides and ORFs), two cell types (U2OS and A549) and two time points per perturbation modality (Supplementary Table 1). This UMAP plot includes the CPJUMP1 primary experiment (4 Compound, 4 CRISPR and 2 ORF plates per cell type and time point) plus all other data points from the CPJUMP1 experiments, as outlined in Figure 3.

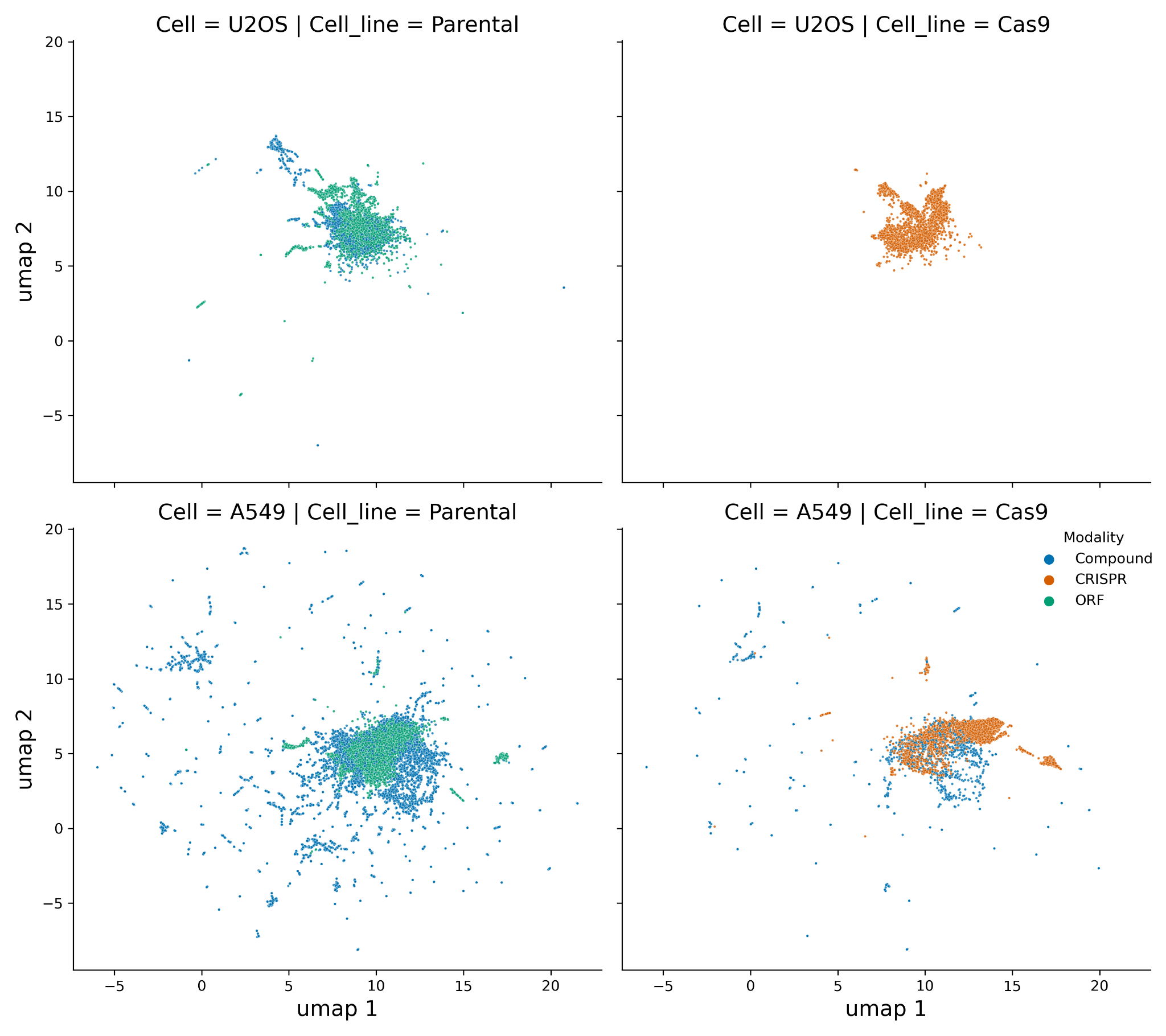

Supplementary Figure 7: **Cas9 status**. Parental line is the original cell line with no modifications. Cas9 cell line is a polyclonal cell line expressing Cas9 (used for all CRISPR and one compound experiment). This UMAP plot includes the CPJUMP1 primary experiment (4 Compound, 4 CRISPR and 2 ORF plates per cell type and time point) plus all other data points from the CPJUMP1 experiments, as outlined in Figure 3.

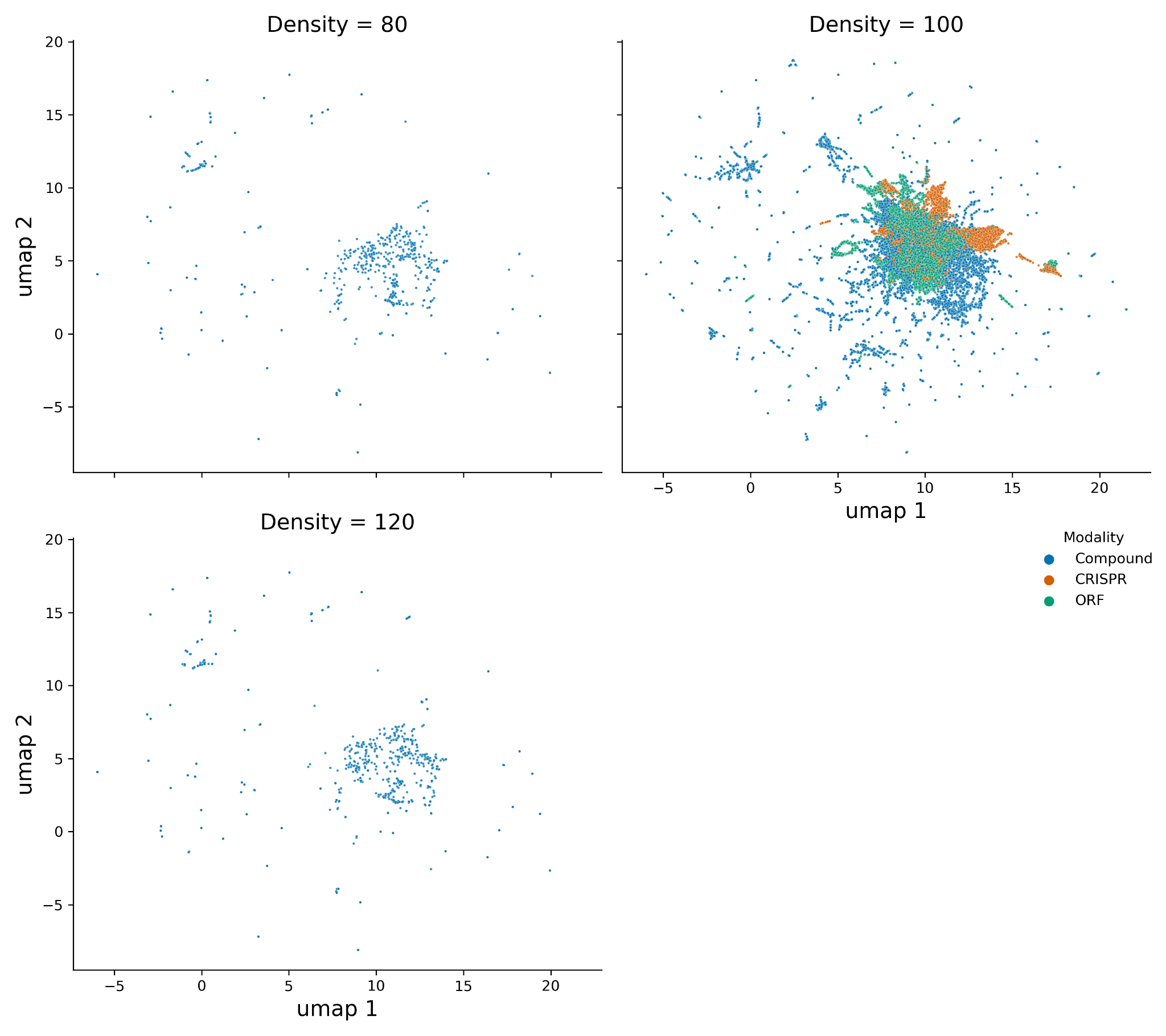

Supplementary Figure 8: **Different cell seeding densities**. Experiments were performed with the baseline seeding density (100%; 1000 cells/well), increased seeding density (120%), and decreased seeding density (80%). This UMAP plot includes the CPJUMP1 primary experiment (4 Compound, 4 CRISPR and 2 ORF plates per cell type and time point) plus all other data points from the CPJUMP1 experiments, as outlined in Figure 3.

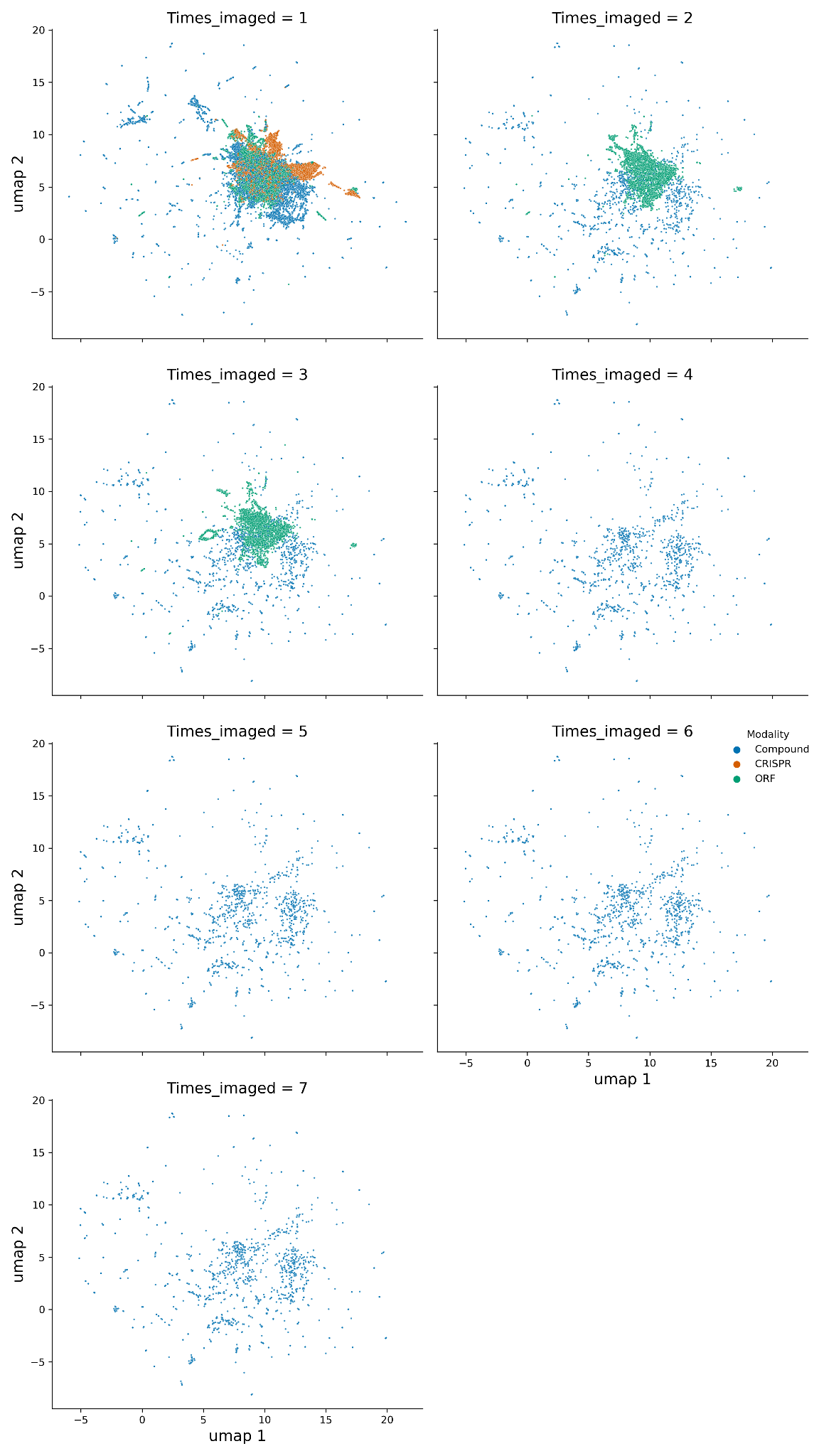

Supplementary Figure 9: **Impact of repeat imaging**. Some plates were imaged more than once. This UMAP plot includes the CPJUMP1 primary experiment (4 Compound, 4 CRISPR and 2 ORF plates per cell type and time point) plus all other data points from the CPJUMP1 experiments, as outlined in Figure 3.

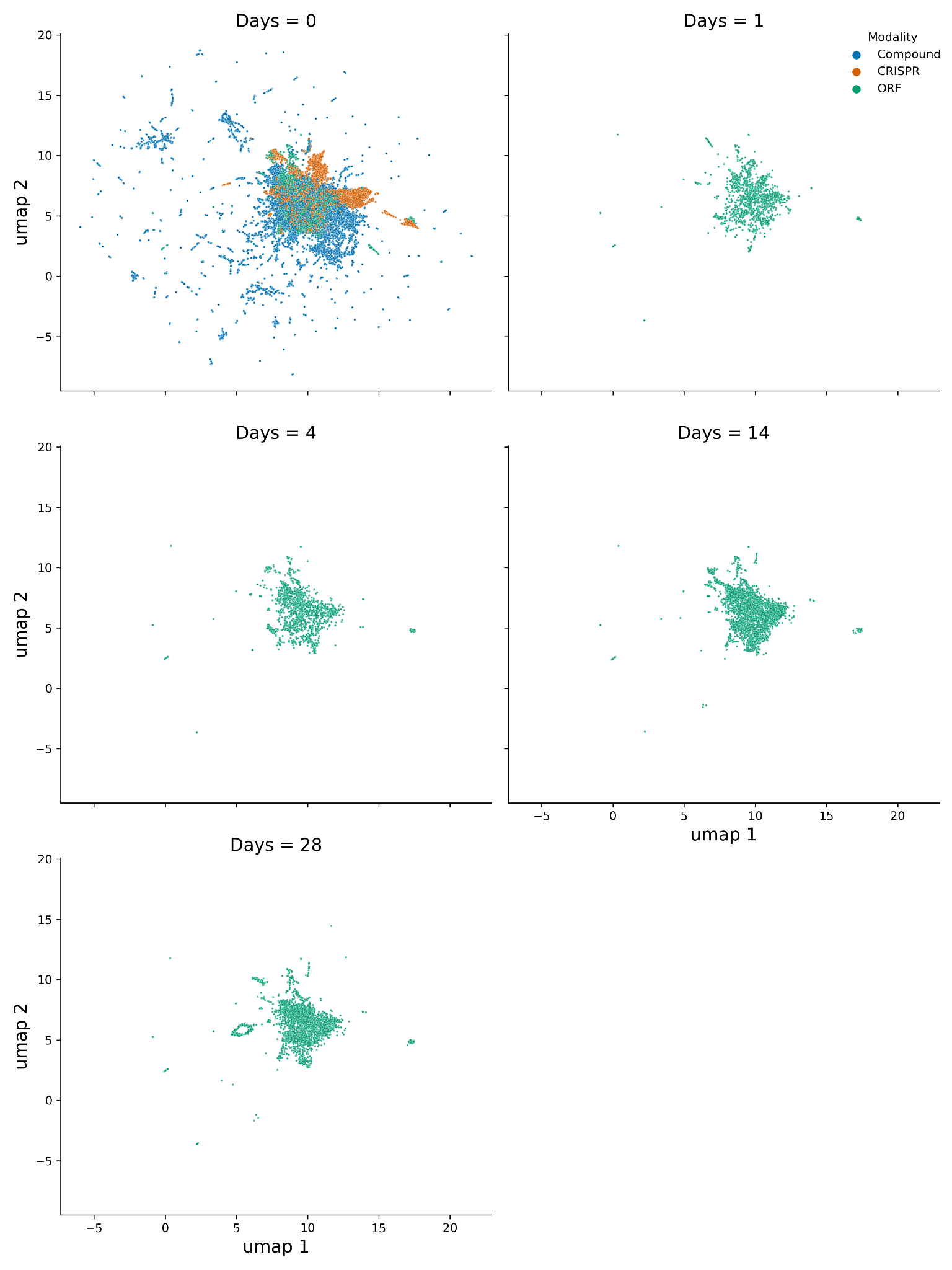

Supplementary Figure 10: **Imaging after a time delay**. A subset of plates were imaged after a certain number of days. This UMAP plot includes the CPJUMP1 primary experiment (4 Compound, 4 CRISPR and 2 ORF plates per cell type and time point) plus all other data points from the CPJUMP1 experiments, as outlined in Figure 3.

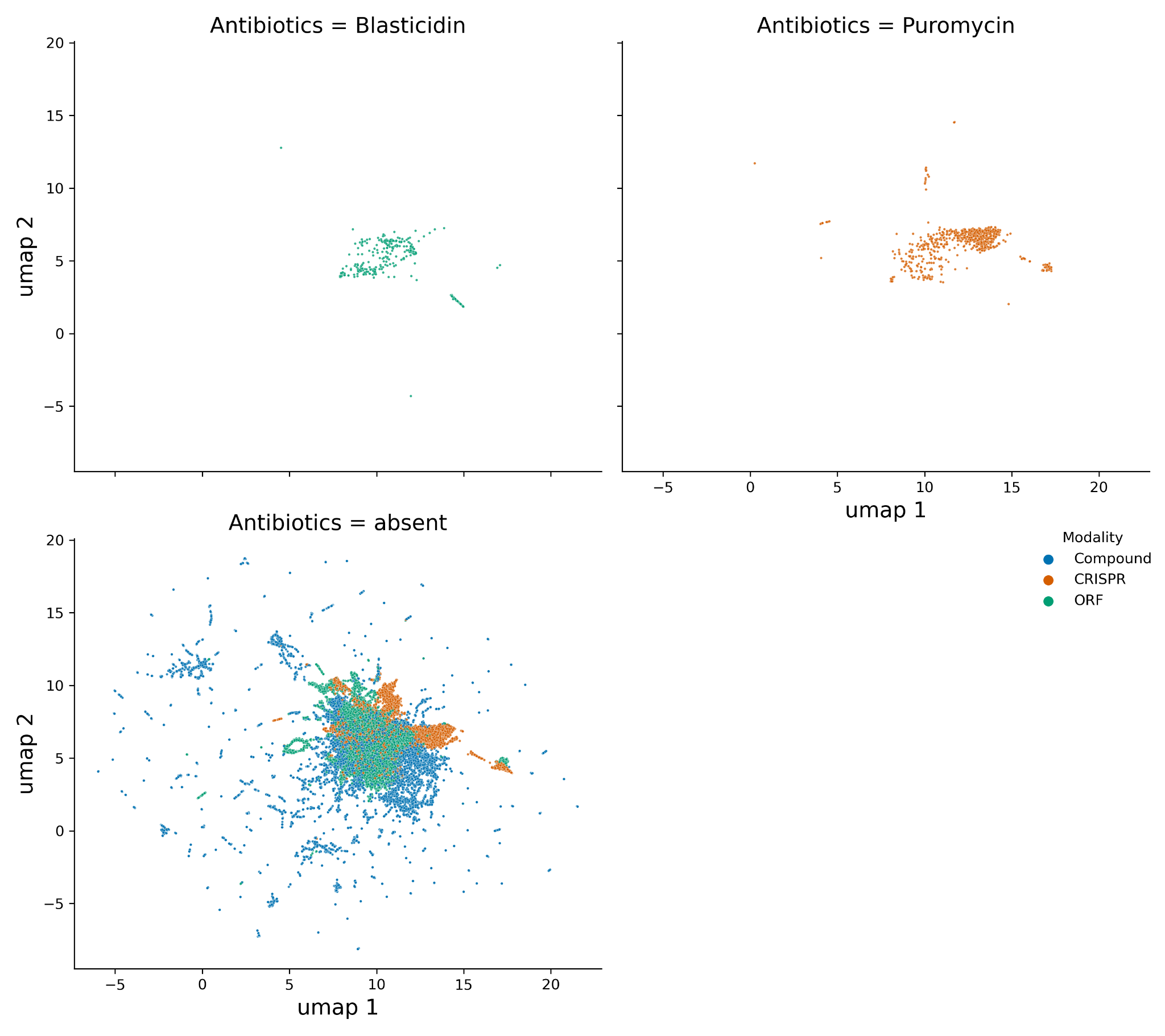

Supplementary Figure 11: **Antibiotic selection**. In some CRISPR and ORF plates, cells were selected using antibiotics. This UMAP plot includes the CPJUMP1 primary experiment (4 Compound, 4 CRISPR and 2 ORF plates per cell type and time point) plus all other data points from the CPJUMP1 experiments, as outlined in Figure 3.
